# Morula complementation restores fetal kidneys in xenocompatible *SALL1* null sheep

**DOI:** 10.64898/2026.07.17.739226

**Authors:** Sarah J Appleby, Lisanne M Fermin, Stephanie Delaney, Jingwei Wei, Fanli Meng, Pavla Turner, David N Wells, Alan J Davidson, Björn Oback

**Author notes:** **SJA**: Department of Clinical and Biomedical Sciences, Faculty of Health and Life Sciences, University of Exeter, RILD Building, Barrack Road, Exeter, EX2 5DW, United Kingdom. **LMF**: Tāwharau Ora- School of Veterinary Sciences, Massey University, Palmerston North, New Zealand. **FM**: Bioeconomy Science Institute, Tuhiraki, 19 Ellesmere Junction Road, Lincoln 7608, New Zealand. **DNW**: 95 Huntington Drive, Hamilton 3210, New Zealand. **BO**: Bioeconomy Science Institute, Nelson Research Centre, 297 Akersten St, Nelson 7010, New Zealand.

## Abstract

To meet the global shortage of organs, extensive genome modifications have been performed to “humanize” livestock tissues for xenotransplantation. However, residual immune rejection remains a problem, prompting alternative approaches that explore the use of animals as hosts for growing human organs. This requires genome editing to disable organogenesis in the host and embryo complementation with suitable donor cells to fill the empty organ niche in chimaeric animals. Pigs have been the predominant livestock species investigated for this approach. Here, we used domestic sheep as hosts for donor-derived kidney formation. Spalt-Like Transcription Factor 1 (*SALL1*) was targeted in male fibroblasts lacking xenoantigens *CMAH* and *GGTA1,* using either one gRNA within zinc finger cluster (ZFC) 2 or two gRNAs to remove all ZF domains. Following somatic cell cloning and embryo transfer of triple knockout strains, fetuses were collected on gestational day 48 to analyze the *SALL1* KO phenotypes. Single gRNA editing produced a hypomorph with different degrees of metanephric hypoplasia, while the dual-gRNA deletion resulted in a null allele which completely abolished nephrogenesis. Female donor cells carrying high vs low copy numbers of an mCherry transgene, as well as *CMAH* and *GGTA1* edits, were used for morula complementation. Fetal kidney development was anatomically and histologically restored in sex-chimaeric hosts, providing proof-of-concept for using sheep as a new model species for in vivo organ generation.

## Introduction

The global gap between short supply and high demand of donor organs could be potentially closed by using genetically engineered livestock for xenotransplantation. Extensive efforts have gone into generating “xenocompatible” livestock species by removing major xenoantigens, such as galactose- α1,3-galactose (α-Gal) (Galili, 2015; Tector et al., 2020), sialic acid *N*-glycolylneuraminic acid (Neu5Gc) (Zhu & Hurst, 2002), and DBA-reactive glycans (or Sd(a) antigen) (Byrne et al., 2014).

Following transplantation into baboons, hearts and kidneys from α-Gal negative pigs survived up to 179 days (Tseng et al., 2005) and 16 days (G. Chen et al., 2005), respectively. Removal of both α-Gal and Neu5Gc in double knockout (DKO) pigs lacking the enzymes α1,3-galactosyl transferase 1 (GGTA1) and cytidine monophosphate-Neu5Ac hydroxylase (CMAH), respectively, lowered the immune response from human sera compared to *GGTA1* KO alone (Lutz et al., 2013). Further study showed that the human immune response elicited by *GGTA1*/*CMAH* KO pig serum was also lower than that elicited by serum from chimpanzee, the species with greatest xenotransplant survival in early studies (Burlak et al., 2014). Continued optimization of pig cells for xenotransplantation entailed further genetic modifications, including knockouts of *CD46* (Lee et al., 2016), *β4GalNT2* (Estrada et al., 2015), and/or *CIITA* (Xu et al., 2024), as well as human transgene knock-ins (Griffith et al., 2022). These multi-gene modifications culminated in the first pig-to-human heart transplant using a donor pig with ten gene edits, including four pig gene KOs and six human transgene knock-ins to render the cardiac graft more xenocompatible (Griffith et al., 2022). More recently, transplantation of a porcine kidney with 69 genomic edits, including inactivation of porcine endogenous retroviruses, and insertion of seven human transgenes, resulted in sustained kidney function without evident xenograft rejection, albeit under expanded immunosuppression (Kawai et al., 2025). These models now define the gold standard for clinical transplants, achieving unprecedented survival times in living patients.

Nevertheless, residual immune rejection, physiological host-donor incompatibilities and potential transmission of animal pathogens pose significant challenges to the widespread adoption of genetically modified livestock organs. To circumvent these issues, new technologies are moving towards using livestock as hosts for growing human organs. Suitable hosts may provide correct in vivo development and function of human organs, without the issues associated with generating complex organ structures in vitro. Generating human organs within a non-human species is an ambitious concept. In practice, it requires the combination of gene editing to ablate the target organ in the host and complementation with heterologous cells of a different genetic origin to restore the organ in chimaeric animals (Bigliardi et al., 2025). As a first proof-of-principle, wild-type (WT) cells were injected into an early blastocyst embryo that had been edited to prevent development of lymphocyte lineages (J. Chen et al., 1993). As the edited “host” was unable to form these cells, complementing WT “donor” cells generated B and T lymphocytes of a single genetic origin in chimaeric mice (Usui et al., 2012). Similar embryo complementation studies were conducted with germline-ablated host and donor mice (Koentgen et al., 2016) and sheep (McLean et al., 2025), even enabling successful germline restoration between closely related rodent species (Kobayashi et al., 2021; Zvick et al., 2022). This “open organ niche” concept was validated in whole organs, producing heterologous liver (Espejel et al., 2010), pancreas (Kobayashi et al., 2010), kidney (Espejel et al., 2010; Usui et al., 2012), lung (Kitahara et al., 2020; Miura et al., 2023; Mori et al., 2019), thyroid (Q. Ran et al., 2020; Wen et al., 2021; Yamazaki et al., 2022), salivary glands (Tanaka et al., 2024), and heart and vascular system (X. Wang et al., 2020) in intraspecific rodent chimaeras. Interspecific complementation successfully generated mouse-rat chimaeras with restored pancreas (Kobayashi et al., 2010), kidney (Goto et al., 2019), lung (Wen et al., 2024), and heart and vascular system (X. Wang et al., 2020). Thus, species with differently sized organs were able to adapt to the size of the host embryo. Intraspecific heterologous complementation has also been demonstrated in non-rodent species via blastocyst complementation in pig for pancreas (Matsunari et al., 2020), liver (Matsunari et al., 2020; Ruiz-Estevez et al., 2021), and kidney (Matsunari et al., 2020; J. Wang et al., 2023). To date, no whole organ niche complementation work has been reported in clinically relevant large animal species other than pig. Kidney is the most needed organ globally. As kidney failure occurs worldwide, kidneys represent the most transplants each year, accounting for over 63% of all solid organ transplants in 2024 (Martin et al., 2026). It is also in highest demand, with about 90% of patients on organ transplant waiting lists specifically waiting for a kidney across many countries (Martin et al., 2026). Since modern kidney transplants have excellent outcomes, with 1-year patient survival rates often exceeding 95%, transplantation is widely considered the most effective and cost-efficient treatment for end-stage renal disease compared to lifelong dialysis (Martin et al., 2026). Unlike heart or liver failure, which are typically fatal without a transplant, patients with kidney failure can be kept alive for years using dialysis, which creates a growing pool of candidates in waiting. Kidney development is conserved across mammals and proceeds via three sequential stages: the pronephros, mesonephros, and metanephros. As development progresses, each increasingly complex renal structure forms while the preceding structure regresses. In mammals, the pronephros is non-functional, whereas the mesonephros can serve as a transient excretory organ during embryogenesis. The definitive kidney, the metanephros, arises through reciprocal interactions between the ureteric bud and the metanephric mesenchyme. These interactions drive ureteric bud branching and mesenchymal differentiation, ultimately giving rise to the collecting duct system, nephrons, glomeruli, and renal stroma (Cullen-McEwen et al., 2016). The coordinated development of these structures is regulated by a network of transcription factors, among which Spalt-like transcription factor 1 (SALL1) plays a central role in establishing and maintaining the metanephric mesenchyme interaction (Nishinakamura et al., 2001).

Here, we used Cas9-mediated genome editing to disrupt both copies of *SALL1*, *CMAH,* and *GGTA1*, simultaneously in ovine fetal fibroblasts. Edited triple KO (TKO) donor cell strains were used to clone fetuses that displayed hypomorphic or no kidneys, depending on the edit. Following complementation of male TKO hosts with female *SALL1* WT donors, kidneys were restored in fetal chimaeras, indicating that the empty kidney niche can be successfully repopulated. This shows that the molecular mechanisms of kidney development and compensation are conserved between rodent and ruminants, supporting sheep as a useful model for xenotransplantation.

## Materials and Methods

Experiments complied with the New Zealand Animal Welfare Act 1999 and were approved by the Ruakura Animal Ethics Committee (applications 14641 and 15078).

### Cell source and culture

Ovine fetal fibroblasts (OFFs) were previously isolated from male (OFF3) and female (OFF4) abattoir fetuses using the hanging drop method, a standard operating procedure in cattle (Appleby et al., 2026). This method was also used to rejuvenate the *CMAH, GGTA1,* and *SALL1* (*CGS*) TKO cell line from a fetus recovered at day 77. Cells were cultured in Dulbecco’s Modified Eagle Medium: Nutrient Mixture F-12 with GlutaMAX™ supplement (DMEM/F12; Gibco, Waltham, MA, USA) with 10% fetal bovine serum (FBS; Moregate Biotech, New Zealand) at 38°C in a humidified 5% CO_2_ incubator. Passaging was performed using a 0.25% trypsin-EDTA (Gibco) incubation for 2–5 min and centrifugation at 200 x *g* for 3–5 min.

### DNA extraction

Genomic DNA was isolated from cells by incubating in lysis buffer (100 mM of Tris [pH 8], 1 mM of EDTA, 0.5% (v/v) of Tween-20, and 0.5% (v/v) of Triton X-100) containing 1 mg/mL of Proteinase K (QIAGEN, Germany) at 55°C for 15 min, followed by a 5 min incubation at 95°C to heat inactivate the Proteinase K. DNA was either ethanol precipitated to obtain a purified sample or used as a crude lysate.

Genomic DNA was extracted from snap frozen tissue samples using DNeasy Blood and Tissue kit (QIAGEN) according to the manufacturer’s instructions. Frozen tissue was digested, and additional steps of RNase A incubation and second final elution were performed.

### Endpoint polymerase chain reaction (PCR)

Endpoint PCR was used to amplify genomic regions of interest using KAPA2G (Kapa Biosystems, Wilmington, MA, USA) following the kit instructions. The reaction contained 0.5 µM of primers and 1 µL of template DNA (total volume of 25 µL). Cycling conditions were as follows: initial denaturation at 95°C for 3 min; 35 cycles of 95°C for 15 s, 60°C for 15 s, and 72°C for 1 s; and final extension at 72°C for 5 min. Amplification of the target region was confirmed on a 1% agarose gel with 1 Kb Plus DNA ladder (Invitrogen, Waltham, MA, USA). PrimerBLAST (Ye et al., 2012) was used to design primers (Table S1) with the parameters set to species: *Ovis aries* and product size: 400–1000 bp. Generally, primers were designed with the region of interest at least 50 bp away from one primer, which would then be used for sequencing.

#### High fidelity PCR

Endpoint PCR with a high-fidelity polymerase was used when analyzing large amplicons. PCR was performed using the KAPA HiFi HotStart PCR Kit (Kapa Biosystems) according to the manufacturer’s instructions. Cycling conditions were as follows: initial denaturation at 95°C for 3 min; 30 cycles of 98°C for 20 s, 60°C for 15 s, and 72°C for 111 s (30 s/kb); and final extension at 72°C for 5 min.

### Sanger sequencing

Amplified DNA regions were purified from the PCR reaction using NucleoSpin Gel and PCR Clean-up kit (MACHEREY-NAGEL, Germany), according to the manufacturer’s instructions. DNA concentration was quantified using a Nanodrop (version 1000; Thermo Fisher, Waltham, MA, USA). Sanger sequencing was performed by Massey University Genome Service (Massey University, New Zealand). Geneious Prime (Biomatters, New Zealand) was used to map the sequences to the reference genome and generate a consensus sequence.

### Droplet digital PCR (ddPCR)

The number of *mCherry* insertions into the sheep genome was estimated using a ddPCR copy number variation (CNV) assay. Beta-casein (*CSN2*) was used as a reference gene (two copies per diploid genome) to measure the ratio of *mCherry* copies using a previously designed assay (McLean et al., 2025). ddPCR was carried out using the QX200 ddPCR system (Bio-Rad, Hercules, CA, USA), according to the manufacturer’s instructions. To separate any tandem insertions, DNA was digested with *Hind*III (Roche, Switzerland) in Roche SuRe/Cut B buffer for 1 h at 37°C. Reactions typically contained 25–70 ng of DNA, 900 nM of primers, and 250 nM of probes (20X stock made in TE buffer: 10 mM of Tris, 0.1 mM of EDTA, pH 8) in ddPCR™ Supermix for Probes (No dUTP) made up to 22 µL with H_2_O. Droplets were prepared using Droplet Generation Oil for Probes. PCR was performed on the C1000 Touch Thermal Cycler (Bio-Rad) with the following cycling conditions: initial enzyme activation 95°C for 10 min; 40 cycles of 94°C for 30 s, and 60°C for 1 min; final enzyme deactivation 98°C for 10 min. Runs were considered successful if the number of generated droplets was >10,000 and positive droplets for HEX (*CSN2* reference gene) was >10. Runs were analyzed on QuantaSoft™ Analysis Pro (Bio-Rad) and the threshold for positive droplets was manually set based on the results of positive and negative template controls run within each experiment.

Cell samples from isolated strains were run in triplicate and controls in duplicate. *mCherry* negative controls included OFF4-derived *CMAH*^−/-^ *GGTA1*^−/-^ strain 56 (#56) and OFF4 WT (Appleby et al., 2026). Two *mCherry* positive controls included OFF3 strains containing *mCherry* insertions of 3 and 8–9 copies, previously determined with this assay (#5 and #9, respectively; (McLean, 2019)). Single replicates of WT male OFF3 (*mCherry* negative) control and no template control (NTC; *mCherry* and *CSN2* negative) were also included.

### Genome editing

#### gRNA-Cas9 guide design for *SALL1*

Previous targeting sites in pig were used to inform target design in sheep, along with the potential to disrupt the majority of the functional domains. Similarity of the parental OFF3 cell line to the reference sheep genome was confirmed for the target regions of *SALL1* using Sanger sequencing.

Following sequence confirmation, constructs were designed for two different targeting strategies. The first strategy was to introduce indels in the middle of exon 2. Three gRNA were designed (gRNA1, gRNA2, and gRNA3; Sequences in Table S1) on CRISPOR (Haeussler et al., 2016) to target 5’ of the C2H2 zinc finger (ZF) domain 5 within ZF cluster 2 (ZFC2). The second editing strategy was to generate a large deletion, >3 kb, removing nearly all of the functional domains within exon 2. Guides were designed to target as close as possible to the 5’ and 3’ ends of exon 2 (gRNA4 and gRNA5, respectively). Targeting constructs were chosen that had high on-target efficiency, low off-target sites, and were near the desired location. Guide sequences beginning with “A”, “T”, or “C” had a single “G” nucleotide added to the 5’ end to improve transcription from the U6 promoter.

Complementary oligos for the top and bottom strands of the gRNA sequence were ordered with appropriate overhangs for ligation into Cas9 plasmids (F. A. Ran et al., 2013).

#### Plasmid generation

To allow for selection of multiple plasmids simultaneously, the Cas9 plasmid PX459 (gifted from Feng Zhang; Addgene plasmid 62988) was modified to have hygromycin resistance (PX459-Hygro; Supplementary methods). *SALL1* gRNA1, 2, 3, and 5 were inserted into PX459-Hygro and gRNA4 was inserted into PX459 following the protocol from the Zhang lab (F. A. Ran et al., 2013). Insertion of the single gRNA into the plasmid was confirmed by performing endpoint PCR using the top gRNA oligo as the forward primer and a reverse primer binding within the CBh promoter. Following band confirmation on an agarose gel, the plasmid sequence was confirmed by Sanger sequencing using a primer upstream of the gRNA insertion site.

#### Transfection and editing efficiency

*SALL1* gRNAs, alongside *CMAH* and *GGTA1* gRNA designed previously (Appleby et al., 2026), were transfected into OFF3 cells. Additionally, mCherry reporter OFF4 cell lines were generated for use in chimaera experiments. These OFF4 cells were transfected with an mCherry plasmid driven by a CAGGS promoter (McLean et al., 2025), *Sleeping Beauty* (SB100) transposase (gifted from W. Kues (Mátés et al., 2009)), and *CMAH* and *GGTA1* gRNAs. All transfections were completed using the Neon® Transfection System (Thermo Fisher), according to the manufacturer’s instructions, with 1.5 × 10^5^ cells and the 10-µL electrode pipette tip. The concentration and volume of all plasmids did not exceed 1.65 µg of DNA per 1 × 10^5^ cells and 10% of the total reaction volume. Cells were electroporated at 1,500 V with one 20 ms pulse and seeded into a prepared 4-well or 35-mm tissue culture plate. Transient selection for 48 h with hygromycin (200 µg/mL) was used to enrich for PX459-Hygro and puromycin (2 µg/mL) for PX459 or *mCherry* plasmid uptake. Cells were grown for 7 d before genomic DNA was isolated.

Editing efficiency of small indels was measured using the Tracking of Indels by DEcomposition (TIDE) webtool (Brinkman et al., 2014). PCR-amplified genomic DNA from the target region in transfected cell samples was Sanger sequenced and sequencing files (abi) were uploaded to the TIDE website.

The indel size setting was adjusted to 35 for all gRNA. All other settings were left at default for *SALL1* and *GGTA1* gRNA. The left boundary window was adjusted to 70 for *CMAH* gRNA to improve the fit score. Results were accepted if the *R^2^*value was ≥0.79.

To evaluate the efficiency of generating a large deletion with two gRNAs, high fidelity PCR was performed on the entire length of exon 2. The KAPA HiFi polymerase was used instead of standard polymerase to amplify both the WT amplicon (3.7 kb) as well as any large deletions (≤1 kb). A gBlock® Gene Fragment (Integrated DNA Technologies, Coralville, IA, USA) was designed to mimic a large deletion, generating a 446 bp amplicon.

*mCherry* insertion efficiency was determined through visualization of mCherry fluorescence on an EVOS microscope (Thermo Fisher).

#### Strain isolation and screening

Clonal edited cells strains were isolated using manual selection of mitotic doublets into a 96-well plate, as described previously (McLean et al., 2021). For cells transfected with the *mCherry* plasmid, only cells with visible fluorescence were passaged. Cells were expanded through 48- and 24-well plates, before cryopreservation, DNA isolation, and endpoint PCR. Editing events that could be resolved as two individual bands were excised from the gel and DNA purified using the NucleoSpin kit. Alternatively, DNA was purified directly from the PCR mixture if two bands were not visualized.

Strains were sequenced and analyzed using Geneious and TIDE to identify desired edits. Strains were considered clonal if there was a maximum of two edits at a single locus (one edit per allele). “Mutation profiles” were used to group strains with identical indels, indicating they likely arose from the same original edited cell. Desirable mutation profiles for small editing events (*SALL1*, *CMAH*, and *GGTA1*) contained indels that caused a frameshift (indel not a multiple of 3) in all genes targeted. For the large *SALL1* deletion, the desired mutation was >2 kb deletion occurring within exon 2 (deletion not extending into the intron). Once strains were selected, PCR products were sub-cloned into bacteria to identify the exact position of the edit identified by TIDE.

Selected strains were assessed for integration of the Cas9 plasmid sequence from PX459 or PX459- Hygro into the sheep genome. Primers were designed to bind within the Cas9 coding sequence, endpoint PCR was performed, and Cas9 presence was confirmed by a band on an agarose gel. PX459 and WT (non-transfected) DNA were used as positive and negative controls, respectively. A sequence known to amplify reliably (*CMAH*, *GGTA1,* or gRNA4 off-target site 1) was run alongside the Cas9 insertion assay to confirm DNA was present from each sample. Strains positive for Cas9 were excluded from further analysis.

Selected strains were then assessed for off-target effects using a biased screen. The top three off-target sites were identified using CRISPOR for each gRNA (*CMAH*, *GGTA1*, and *SALL1* gRNA2, gRNA4, and gRNA5). For *SALL1* gRNA2, the WT sequence of the parental OFF3 and OFF4 cells did not match the reference genome for the third ranked site and the next highest-ranked target site was used. Off-target sites were amplified, sequenced, and compared with the WT sequence for any evidence of editing.

Two TKO strains were selected for further work. Nomenclature used for the TKO strains is based on the strain number and kidney KO phenotype: the small indel strain is referred to as *CGS4^hypo^* (*CMAH^− 34/-44^ GGTA1^+1/-303^ SALL1^+1/-11^* strain #4 with a hypomorphic phenotype) and the large deletion strain is *CGS20^null^* (*CMAH^−34/-44^ GGTA1^+1/-303^ SALL1^Δ32741/Δ32741^* strain #20 with a null phenotype). The two mCherry reporter cell strains selected for further work are referred to by their strain number: *mCherry38* (*CMAH*^−1/-19^ *GGTA1*^−3/-3^ mCherry^+^) and *mCherry54* (*CMAH*^−1/-11^ *GGTA1^WT^*^/-4^ mCherry^+^).

### Somatic cell transfer (SCT) cloning and embryo transfer (ET)

Sheep SCT cloning and ET were carried out as described (Appleby et al., 2026). Morula aggregation was used for all cloning runs; aggregation of two embryos from the same cell strain was used to improve embryo quality and in vivo survival, whereas aggregation of embryos from different cell strains was used to generate chimaeras. Cell strains combined to generate chimaeras included *CGS4^hypo^* with *mCherry38* (*CGS4^hypo^*↔*mCherry38*) and *mCherry54* (*CGS4^hypo^*↔*mCherry54*) or *CGS20^null^*with *mCherry54* (*CGS20^null^*↔*mCherry54*; no *CGS20^null^*↔*mCherry38* chimaeras were generated). Blastocysts were morphologically graded on day 6 or 7 and transferred into synchronized recipient ewes or vitrified for future transfer (Robertson & Nelson, 2011). Groups of 3–4 embryos were laparoscopically transferred to recipient ewes; 10 ewes were used per single cell group and 15–20 ewes per chimaera group. Pregnancies were monitored using ultrasound with the M-turbo ultrasound system (SonoSite, Bothell, WA, USA). Between day of gestation (D) 30–40, pregnancy establishment was determined using transrectal ultrasound (L52x, 10-5MHz transducer). Scans after D45 were completed using transabdominal ultrasound (C60x, 5-2MHz transducer). Fetuses were collected on D48 by slaughter recovery and the whole reproductive tract was removed.

### Fetal recoveries

After retrieving the fetus from the reproductive tract, images were taken and weight measured. Fetuses were dissected by first removing the head and forelimbs, followed by a vertical incision through the skin down the ventral midsagittal line. Skin was carefully separated to allow opening of the body cavity and removal of organs, leaving the kidneys attached to the dorsal side of the cavity. The fetus was then moved onto a wax tray and pinned into place for imaging and further dissection. Using microdissection tools under a stereomicroscope, attached metanephros, mesonephros, ureters, bladder, gonads, and adrenals were separated from the body wall and placed in a Petri dish containing phosphate buffered saline (PBS). Mesonephros, gonads, and adrenal were removed to isolate the metanephros, ureters, and bladder, which were imaged and samples taken.

#### Tissue collection

Fetal tissue samples were collected for genotype and phenotype analysis. Samples were collected from skin, muscle, heart, liver, kidney, brain, adrenal, and gonad. In some instances, degeneration of the organ was too great to collect samples from. Collected tissue was dissected into small fragments and snap frozen in liquid nitrogen. Samples were transferred into a single 1.5-mL Eppendorf tube per tissue type and stored at -80°C.

Fixation in PFA was used for histological evaluation of fetal organs. The metanephros, adrenal, and gonad attached to mesonephros from one side of the fetus were isolated and kept whole. The entire back section with kidneys still attached was fixed from fetuses in the late stages of degeneration. Samples were incubated in freshly depolymerized (56°C water bath) 4% (w/v) paraformaldehyde (PFA) with 4% (w/v) sucrose in PBS overnight at 4°C. Samples were then stored in PBS with a small amount of PFA at 4°C until ready for sectioning.

#### Histology

PFA fixed tissue samples were transferred to 70% ethanol and incubated at 4°C overnight. On the following day, tissues were processed through standard dehydration series for paraffin embedding. Tissue was incubated in 1:1 xylol:molten paraffin at 60°C overnight, followed by 100% molten paraffin for 2 days, changing paraffin twice each day. Samples were embedded in paraffin and sectioned. A microtome was used to generate sections of 6 µm for analysis. Sections were transferred onto charged glass slides using a water bath and left to air-dry overnight at room temperature. Slides were stored at room temperature until staining. Hematoxylin and eosin (H&E) staining was performed on paraffin sections using standard protocols. Stained sections were mounted in DPX Mountant for histology (Sigma-Aldrich, St. Louis, MO, USA) under a glass coverslip.

#### Chimaerism analysis

Chimaerism was confirmed using PCR with primers designed to identify the male (*CGS4^hypo^* and *CGS20^null^*) and female (*mCherry38* and *mCherry54)* cell strains. Presence of the male strain was confirmed with primers for the Y chromosome gene *DDX3Y*. Presence of the female strain was confirmed with primers designed within the cut site of *CMAH* in the male strains, specifically binding the -1 bp allele of female *mCherry54*, as well as the WT and *mCherry38* -1 bp allele. Male and female assays were run separately to avoid amplification of the male amplicon being affected by presence of the female amplicon, and vice versa. Endpoint PCR reactions contained 6–90 ng/µL of DNA.

Reactions were combined and run on 1% agarose gel to analyze the banding pattern. A second female specific *CMAH* PCR with 5 µl of DNA (40–80 ng) was performed for fetuses with red fluorescent kidneys if no female DNA amplified for the previous assay, alongside a genomic control with standard *CMAH* primers. Sensitivity of the PCR assay was determined using a dilution series.

Simulated chimaeric contributions of 0.1%–50% were made by combining *CGS4^hypo^* and *mCherry54* DNA; varying levels of DNA from one sex were combined with 50 ng of DNA from the opposite sex to a total of 100 ng DNA. PCR products were run on a 1% agarose gel to determine when the amplicon bands were no longer visible.

### Imaging

Live images of cells and blastocysts were captured using the EVOS microscope. For imaging with Hoechst, cells were incubated in PBS containing 5 µg/mL of Hoechst for 5 min before imaging. Cells were imaged at 20x magnification with light settings of 70% for brightfield, 30%–40% for Hoechst, and 50%, 60%, or 70% for red fluorescence. Blastocysts were imaged at 20x magnification with light settings of 50% for brightfield and 40% or 60% for red fluorescence.

Color images of fetuses were captured using a Canon DSLR camera with 18–55 mm lens for imaging whole fetuses and the kidney in situ, or a 100 mm macro lens for dissected organs. Two images were captured at each time point using the ChemiDoc (Bio-Rad). Colorimetric image mode was used to take black and white images of the subject, with exposure set on Rapid Auto-exposure. mCherry fluorescence was captured using DyLight 550 settings, with exposure set at 0.2 and 2 s. Brightness of DyLight 550 images was set to the same value using ImageLab (Bio-Rad).

H&E sections were imaged using Olympus BX50 microscope (Olympus, Japan) with 2x and 10x objectives. Images were processed using Ocular software (Photometrics, Tucson, AZ, USA).

#### Image analysis

Organ size and curved crown rump length (CRL) were measured from the colorimetric image using ImageJ (version 1.45s; National Institutes of Health, Bethesda, MD, USA). The scale was set on ImageJ using the ChemiDoc image dimensions (large image = 180 x 144 mm, medium image = 95 x 76 mm). Curved CRL was measured on the whole fetus image using the segmented line tool. Area of metanephros, mesonephros, gonads, and adrenals was measured using the polygon tool to draw an outline around the whole organ. Mesonephros, gonads, and adrenals were measured on the image taken following removal from the body cavity (organs attached to each other by fascia). Metanephros was measured on the fully dissected image (with only ureter and bladder still present). In fetuses where organs were not removed due to degeneration, area was measured on in situ image. For analysis, organ area was normalized on weight (mm^2^/g).

mCherry fluorescence was measured on ImageLab using the Volume Tools function. The subject was outlined using the Freehand tool and fluorescence recorded as the Volume (brightness intensity) divided by Area (mm^2^). Images used for analysis were the same as those used to measure organ size.

### Statistical analysis

The data were processed using Excel (Microsoft Office 365, Version 2008; Microsoft, Redmond, WA, USA) or R and the tidyverse suite of packages (Wickham & Grolemund, 2017). Graphs in R were constructed using ggplot2.

For *SALL1* gRNA1–3 editing efficiency, TIDE analysis was performed on both forward and reverse reactions per gRNA (n= 2) and the mean was determined for each indel. Error bars represent standard error of the mean (SEM). Overall efficiency is displayed as the mean for the two individual efficiencies. *R^2^*is shown as the lowest value, indicating the worst predicted fit of the two reactions. For all other gRNA transfections, TIDE analysis was performed on a single reaction per gRNA, with efficiency and *R^2^* being reported directly from the TIDE prediction. For the ddPCR reaction, triplicate or duplicate wells were merged on QuantaSoft, with values presented for the merged data set. Error bars represent the Poisson distribution generated by QuantaSoft.

Reconstructs generated across SCT runs within a year were pooled. The number of cleaved embryos was normalized to the number of reconstructs placed into in vitro culture (IVC). Total blastocyst development rate was normalized on cleaved embryos. *P* values for SCT and ET were calculated using Fisher’s exact test in Excel. Organs from dissected fetuses were normalized on weight (mm^2^/g).

Total chimaerism efficiency was normalized on total fetuses. The 95% confidence interval (CI) for WT males (OFF3) and females (*mCherry54* and OFF4 WT) is represented as shading on the graph to indicate the expected size for that gender.

## Results

### Kidney ablation

#### Generating TKO ovine fibroblasts

To generate TKO male sheep fibroblasts lacking *SALL1*, as well as the xenoantigen-encoding genes *CMAH* and *GGTA1*, all three genes were targeted simultaneously in WT OFF3 cells. We used two proven gRNAs against *CMAH*/*GGTA1* (Appleby et al., 2026) and five novel gRNAs with minimal predicted off-target sites to target *SALL1* (Table S2). Two consecutive editing strategies were employed (Fig. 1a). First, three *SALL1* gRNAs (gRNA1–3) were selected to introduce random indels before C2H2 ZF domain 5 within ZFC2, resulting in frameshift mutations that produced a prematurely terminated nonsense protein (Fig. 1b). Second, we used rejuvenated cell strain 4 (*CGS4^hypo^*) for re-targeting *SALL1* with another two guides (gRNA4, 5) to delete most of intervening exon 2, introducing a frameshift and premature stop codon that ablates all ZF domains (Fig. 1c). Editing efficiency was measured by TIDE analysis (Fig. S1a-c). The most efficient guide for introducing indels was gRNA2 with 97% of the pooled population predicted to be edited, with 73% of indels due to a single “A” insertion. Dual guide editing to introduce a large deletion was analyzed using endpoint PCR in cell strains isolated from gRNA4/5 transfection (Fig. S2a). Nine strains showed deletion bands (2, 3, 6, 8, 14, 15, 16, 20, 21), seven contained only WT-sized bands at 3.5 kb (5, 10, 11, 12, 13, 17, 18) and six had either no or faint WT bands (1, 4, 7, 23, 24, 26) (Fig. S2b). The nine putative large deletion strains were investigated further with primers binding within the intended deleted sequence. Four strains did not amplify the correct-sized product (2, 3, 14, 20), indicating this region had been lost from the sequence (Fig. S2c). Overall, 4/22 (18%) strains contained large deletions on both alleles. Strains 2 and 14 contained the same -2745 bp edits that retained C2H2 ZF domains 6–9, indicating they likely originated from the same edited cell (Fig. S2d). In strain 20, which still contained the 5’ intron-exon 2 junction, the homozygous 3274 bp deletion caused a frameshift in ZF domain 9. Strain 20 was chosen for further analysis as it had the desired mutation and was the only strain still dividing when the other three strains had already become senescent.

**Fig. 1.**
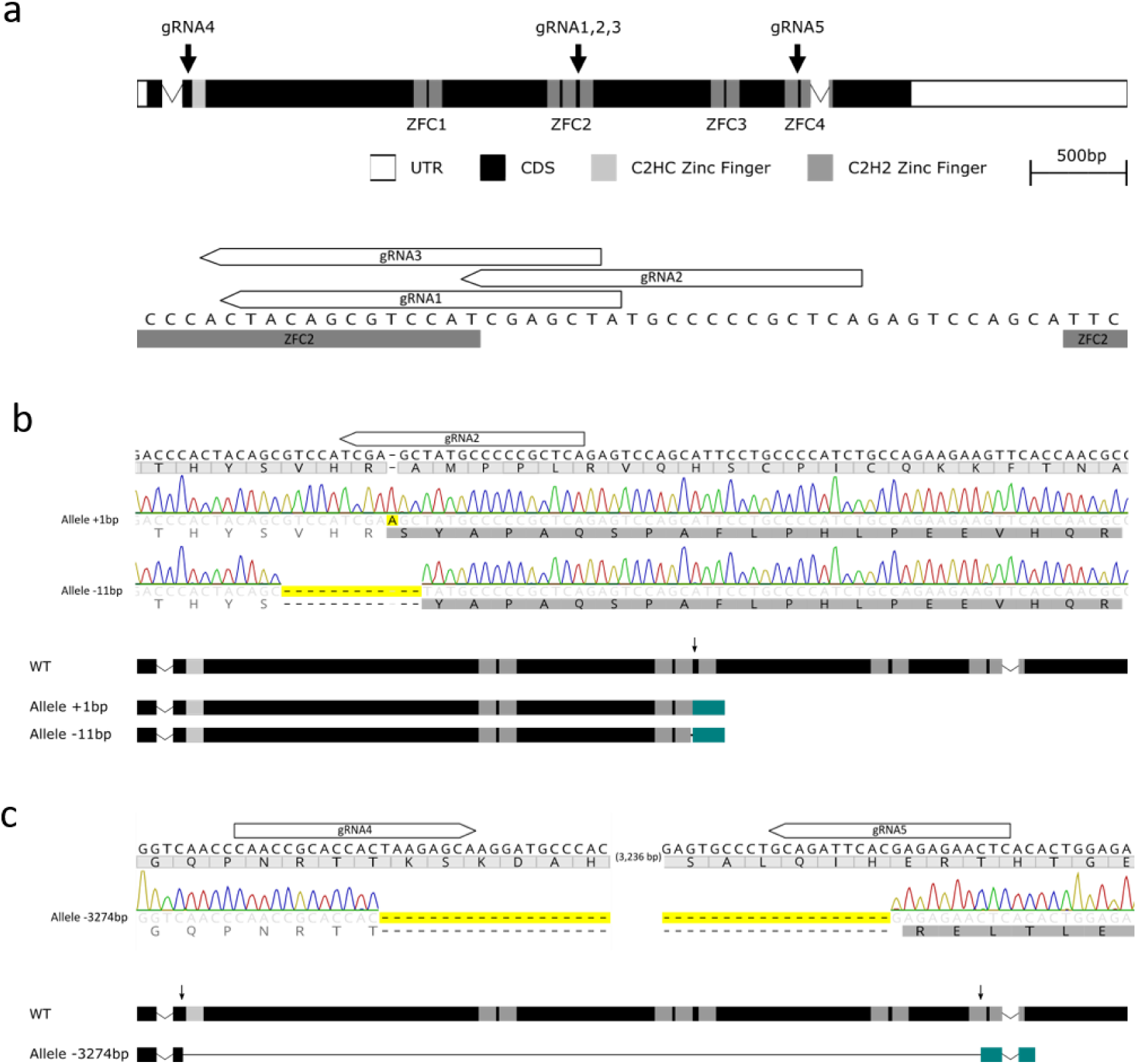
Ovine gene structure, editing locations for *SALL1* and *SALL1-/-* fetal genotype. (a) Gene structure shows the untranslated region (UTR, white), introns (lines), and exon coding sequences (CDS, black) with conserved functional domains (grey): the N-terminal C2HC zinc finger (light grey), followed by nine C2H2 zinc finger motifs organized in four clusters (ZFC1–4, dark grey). Numbered target sites for gRNA editing indicated by arrows. Sequence displayed below shows small editing strategy, along with numbered gRNA binding sites (not including PAM site). *CGS4^hypo^*sequenced for *SALL1* (b) single and (c) dual guide editing strategies. Translation shown under nucleotide sequence. PCR products that could not be separated on agarose gel were sub-cloned into bacteria to sequence individual alleles. Nucleotides sequenced differently from reference shown in black (with changes to amino acid sequence shown underneath in black within shaded boxes), matching nucleotides/amino acids in light grey. Yellow boxes highlight edited nucleotides. Changes to CDS structure shown below sequencing results (black = CDS, grey = zinc finger domains, teal = frame-shifted amino acids).

We PCR-screened TKO strains for unintentional integration of the Cas9 coding region. No Cas9 bands were detected in five small indel TKO strains (4, 5, 8, 19, 21; Fig. S3a, b) and the large deletion strain 20 (Fig. S3c, d). To test specificity of the gRNA-Cas9 system, a biased Sanger sequencing screen of the top three potential off-target sites revealed no mutations in the homozygous male mutant for *CMAH* (Fig. S3e), *GGTA1* (Fig. S3f), *SALL1* gRNA2 (Fig. S3g), gRNA4 (Fig. S3h), and gRNA5 (Fig. S3i). Based on their mutation profile, absence of Cas9 integration and off-target effects, small indel strain 4 (*CGS4^hypo^* = *CMAH^−34/-44^ GGTA1^+1/-303^ SALL1^+1/-11^*) and large deletion strain 20 (*CGS20^null^* = *CMAH^−34/-44^ GGTA1^+1/-303^ SALL1^Δ32741/Δ32741^*) were chosen for SCT (Table S3).

#### Genotype of cloned TKO fetuses

Homozygous male TKO cell strains were chosen to generate fully defined fetal genotypes by somatic cell cloning. Using *CGS4^hypo^*and *CGS20^null^* donor cells, seven SCT runs were completed to generate cloned blastocysts (Table S4). Cleavage of reconstructs was higher for clones generated from the *CGS20^null^* strain (385/428=90%; *p*<0.001) compared to *CGS4^hypo^* (312/409=76%) and WT OFF4 (427/582, 73%; *CGS4^hypo^* vs WT *p*=0.335). Blastocyst development from cleaved embryos was highest for *CGS20^null^*(86/385=22%), followed by *CGS4^hypo^* (50/312=16%; *p*=0.045). Control WT OFF4 had the lowest development (19/427=4%; *p*<0.001 against both groups). High morphological quality blastocysts (Grade 1–2) developed at a similar rate across all groups (52%, 42%, and 37% for *CGS20^null^*, *CGS4^hypo^*, and WT, respectively; *CGS4^hypo^* vs *CGS20^null^ p*=0.325, *CGS4^hypo^* vs WT *p*=0.915, *CGS20^null^* vs WT *p*=0.333).

Cloned blastocysts generated from the three genotypes, as well as male OFF3 WT controls, were transferred into recipient ewes (Table 1). Approximately 30 embryos for each edited and control group were transferred into recipient ewes and established a pregnancy with similar frequency. However, *CGS20^null^* produced the lowest number of fetuses with a heartbeat (1/30=3%; vs *CGS4^hypo^ p*=0.059, vs OFF4 *p*=0.467, vs OFF3 *p*=0.508). Fetuses were recovered on D48 from ewes where a heartbeat was detected. *CGS4^hypo^*and OFF3 matched the scanning prediction, with 4 (13%) and 2 (13%) healthy fetuses recovered, respectively. Two degenerating fetuses were also recovered from *CGS4^hypo^* (6%), one in the early stages of degeneration, indicating that the fetus had recently died, and one that had arrested at approximately D30 (based on CRL). A higher number of healthy fetuses were recovered from OFF4 WT than expected (4/14, 29%) due to the presence of triplets. A single fetus in the early stages of degeneration was recovered from *CGS20^null^* (3%). Overall, OFF4 generated the highest number of healthy fetuses compared to other groups (vs *CGS4^hypo^ p*=0.390, vs OFF3 WT *p*=0.582, vs *CGS20^null^ p*=0.015), and *CGS20^null^*the lowest (vs *CGS4^hypo^ p*=0.121, vs OFF3 WT *p*=0.212; *CGS4^hypo^* vs OFF3 WT *p*=1.00). Fetal DNA was isolated from snap frozen heart samples, and their genotypes were compared to the original edited donor cell. Chromatograms from all fetuses matched that of the original cell chosen for SCT (Fig. S4).

**Table 1.** In vivo survival of TKO cloned host embryos.

| Donor cells | Recipient ewes | Embryos transferred | Scan 1 <D40 |  | D48 fetuses |  |
| --- | --- | --- | --- | --- | --- | --- |
|  |  |  | Pregnant (%)<sup>a</sup> | HB (%)<sup>a</sup> | Collected healthy (%)<sup>a</sup> | Collected aborting (%)<sup>a</sup> |
| OFF3 *CGS4<sup>hypo</sup>* | 10 | 31 | 10 (32) | 7 (23)* | 4 (13)<sup>b</sup> | 2 (6)<sup>c</sup> |
| OFF3 *CGS20<sup>null</sup>* | 10 | 30 | 7 (23) | 1 (3)* | 0 (0)** | 1 (3)<sup>d</sup> |
| OFF4 | 4 | 14 | 4 (29) | 2 (14) | 4 (29)** | 0 (0) |
| OFF3 | 5 | 15 | 4 (27) | 2 (13) | 2 (13) | 0 (0) |
| **Total** | **29** | **90** | **25 (28)** | **12 (13)** | **11 (12)** | **3 (3)** |

<sup>a</sup> Normalized on embryos transferred.
<sup>b</sup> Three fetuses collected on D48 and one fetus collected on D77 that was missed on D48 scanning.
<sup>c</sup> One fetus in early stages of degeneration and one arrested approximately D30.
<sup>d</sup> One fetus in early stages of degeneration.
\* Values within this column differed with $p = 0.059$ .
\*\* Values within this column differed with $p = 0.015$ .
CGS, *CMAH*<sup>-/-</sup> *GGTA1*<sup>-/-</sup> *SALL1*<sup>-/-</sup>; D40, gestational day 40; D48, gestational day 48; HB, heartbeat.

#### Phenotype of cloned TKO fetuses

TKO kidneys from all healthy and early degenerated fetuses displayed variable metanephric kidney phenotypes (Fig. 2a). Some showed either bilateral hypoplasia (F01) or unilateral hypoplasia and agenesis (F02), while others (F07) had two fully formed metanephric kidneys, approximately the size expected from a D48 male fetus (F15). The single fetus from strain *CGS20^null^*that survived to D48 (F21) had no metanephric kidneys. Metanephric kidney histology was qualitatively evaluated on stained sections from one kidney of each fetus (Fig. 2b). The structure of *CGS4^hypo^* kidneys appeared similar to that of WT, showing comparable populations of stromal cells, glomeruli, and tubules, regardless of size. Kidney size ranged from much smaller than expected for a D48 male fetus (F01, F02 = 62%–92% reduced) to moderately smaller (F07, F08 = 20%–40% reduced), but at the upper limits of the observed D48 female fetuses (Fig. 2c). The fetus collected at D77 (F11) also had 2 fully formed kidneys (Fig. S5). However, the left kidney was misplaced, being lower and medially positioned compared to the right kidney. Once dissected out of the body cavity, the ureter of the left kidney could be seen to be positioned cranially and did not connect to the bladder. Kidney hypoplasia or agenesis was observed in 2/5 (40%) *CGS4^hypo^* fetuses collected. By contrast, kidneys were absent in *CGS20^null^* (F21). Within the WT control fetuses, sexual dimorphism of organ size was observed. Mean kidney size was 40% smaller in females compared with males (1.10 mm^2^/g vs 1.80 mm^2^/g).

**Fig. 2.**
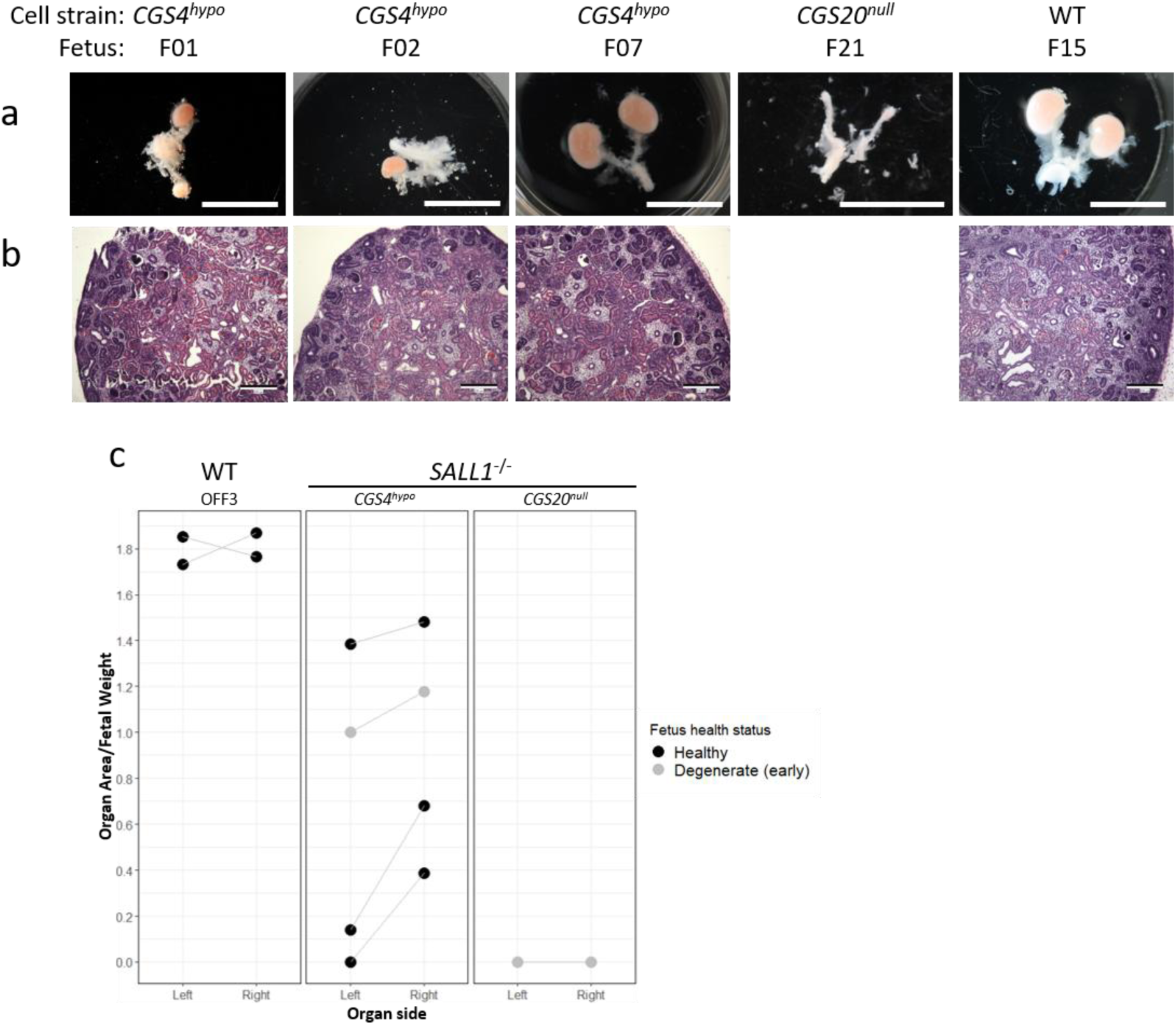
Fetal kidney knockout phenotype. Images from triple knockout (TKO; *CMAH/GGTA1/SALL1 = CGS*) D48 fetuses cloned from cell strains 4 and 20 (*CGS4^hypo^*, *CGS20^null^*) compared to wild-type (WT) showing metanephros (a) morphology (scale bar = 10 mm) and (b) histological hematoxylin/eosin (H&E)-stained sections (scale bar = 200 µm). (c) Size comparison of fetal metanephros area (mm^2^), normalized on fetal weight (g), from TKO fetuses (*CGS4^hypo^*, *CGS20^null^*) vs WT control males (OFF3). Paired organs from a single fetus are connected with grey line.

Apart from displaying a fully anephric phenotype, the surrounding organs (gonads, adrenals, and mesonephros) appeared normal, albeit variable, in size (Table S5). WT adrenals and mesonephroi were similar in both sexes (0.29 mm^2^/g vs 0.36 mm^2^/g, and 0.56 mm^2^/g vs 0.76 mm^2^/g in female and male, respectively), with overlapping 95% CIs. Ovaries were half the size of testes (0.25 mm^2^/g vs 0.48 mm^2^/g).

*CGS4^hypo^* testes were similar to the male OFF3 control, whereas *CGS20^null^* testes were 37% smaller (Fig. S6a). No male OFF3 of similar fetal size to *CGS20^null^* was collected, which may have affected comparison. For both adrenals (Fig. S6b) and mesonephros (Fig. S6c), normalized organ size of the *SALL1* TKO fetuses was within observed ranges for both male and female WT fetuses.

### Donor cells for complementation

### *mCherry* donor cells

To rescue aberrant kidney development in TKO male cloned embryos, we generated genetically traceable female mCherry-transgenic donor cells. The *mCherry* sequence was inserted into OFF4 cells while simultaneously editing *CMAH* and *GGTA1* with the most efficient gRNAs previously identified (Appleby et al., 2026). Strains were sequenced to determine the *CMAH* and *GGTA1* edits, but none showed frameshift mutations in both genes (Table S6). All 18 strains were analyzed for Cas9 integration. One strain was positive for Cas9 sequence inserted into the sheep genome (profile J, strain 46, Fig. S7a, b). Two transgenic clonal strains for SCT were screened for off-target editing.

Strain 38 (*mCherry38*) was initially chosen as it contained a frameshift mutation in *CMAH* and both *GGTA1* alleles were edited (-3 bp). For the lowest number of mCherry insertions, strain 54 (*mCherry54*) was chosen, despite carrying only a heterozygous *GGTA1* mutation (0, -4 bp). No off-target mutations were detected in the top three target sites for *CMAH* (Fig. S7c) and *GGTA1* in either strain (Fig. S7d). The number of *mCherry* insertions in the clonal cell strains was estimated using ddPCR (Fig. 3a). Across all strains, between 2–15 copies of *mCherry* were inserted into the genome with seven strains containing >10 *mCherry* copies (39%), nine containing 5–10 copies (50%), and two containing <5 (11%). Only one strain contained <2 copies (*mCherry54*). Copy number variance within strains of the same mutation profile was generally low (± 1 copy) but increased for strains with higher number of *mCherry* insertions (profiles F 15.5 vs 11.2 copies, and E 13.4 vs 9.0). Red fluorescence intensity for selected strains (*mCherry38*, *mCherry54*) correlated with the estimated number of *mCherry* copies (Fig. 3b). At non-saturating settings for high-copy strain *mCherry38* (50% light), fluorescence from low-copy strain *mCherry54* was hard to detect, while at optimum settings for *mCherry54* (70% light), *mCherry38* was overexposed.

**Fig. 3.**
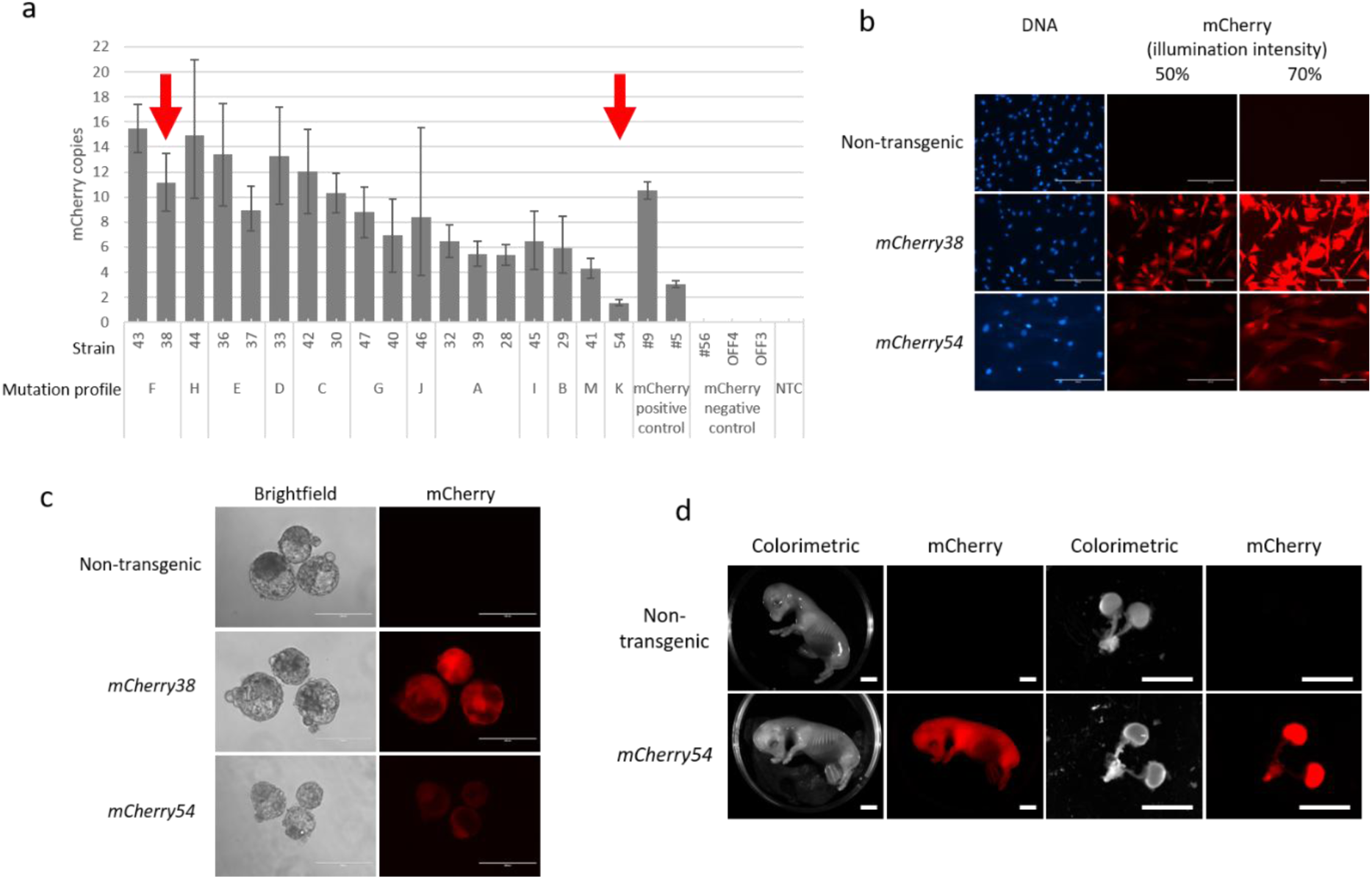
Characterization of mCherry-overexpressing donor cell lines. (a) Droplet digital PCR (ddPCR) quantification of genomic mCherry insertions in clonal OFF strains, normalized on reference gene CSN2. Previously characterized OFF3 mCherry strains (#5, #9) served as positive controls. Non- transgenic OFF4 CMAH^−/-^ GGTA1^−/-^ (#56), parental OFF4, and parental OFF3 cell lines provided negative controls (No template control = NTC). Error bars are the max/min error generated by the QuantaSoft software. OFF4 mCherry strains n=3 technical replicates, and control samples n=2, except OFF3 and NTC (n=1). (b) mCherry fluorescence in clonal OFF4 strains (*mCherry38*, *mCherry54*) compared to non- transgenic parental OFF4 cells. Images show DNA stain (Hoechst 33342) and red fluorescence viewed at 50% and 70% illumination intensity. Scale bar = 200 µm. (c) Red fluorescent signal in cloned blastocysts derived from *mCherry38* and *mCherry54* vs non-transgenic donors (*CGS4^hypo^*). Fluorescent images were captured at 40% light intensity. Scale bar = 200 µm. (d) Representative recovered healthy fetuses on D48, derived from non-transgenic parental OFF4 line without edits or insertions (F14) and *mCherry54* (F13). Colorimetric images show whole fetus and isolated metanephros (ChemiDoc, 0.2 s exposure on DyLight550 channel). Scale bar = 10 mm.

### *mCherry* donor embryos and fetuses

High- and low-copy strains *mCherry38* and *mCherry54*, respectively, were used for SCT cloning (n=6, Fig. 3c, Table S7). Total blastocyst development was the same for *mCherry38* (32/267=12%) and *mCherry54* (71/572=12%) with similar proportions of high morphological grade blastocysts in both groups (16/32=50%, and 38/71=54%, for *mCherry38* and *mCherry54*, respectively). At non-saturation illumination (40% light), mCherry fluorescence intensity from *mCherry38* was about three times higher than *mCherry54* (*n*=3). Blastocysts generated from the two strains were transferred into recipient ewes and recovered on D48 to assess fluorescence in the whole fetus and isolated metanephros (Fig. 3d). Cloned embryos from both donor strains established pregnancies with similar rates, but none of the high-copy *mCherry38* fetuses had a heartbeat (0/27 vs 6/59 for *mCherry38* vs *mCherry54*; *p*=0.192). Pregnancy establishment of *mCherry38* fetuses was significantly lower than for *CGS4^hypo^* controls (*p*=0.02). Two healthy *mCherry54* fetuses were recovered, one of which still had a heartbeat at D48 (Table 2). Sequencing of the edited alleles in heart samples from both fetuses confirmed that they were identical to the cell strain used for SCT (Fig. S8). All edits matched the size identified by TIDE.

**Table 2.** In vivo survival of mCherry cloned donor embryos.

| Donor cells | Recipient ewes | Embryos transferred | Scan 1 <D40 |  | D48 fetuses |  |
| --- | --- | --- | --- | --- | --- | --- |
|  |  |  | Pregnant (%)<sup>a</sup> | HB (%)<sup>a</sup> | Collected healthy (%)<sup>a</sup> | Collected aborting (%)<sup>a</sup> |
| *mCherry38* | 9 | 27 | 7 (26) | 0 (0) | 0 (0) | 0 (0) |
| *mCherry54* | 19 | 59 | 17 (29) | 6 (10) | 2 (3) | 2 (3)<sup>b</sup> |
| **Total** | **28** | **86** | **24 (28)** | **6 (7)** | **2 (2)** | **2 (2)** |

<sup>a</sup> Normalized on blastocysts transferred.
<sup>b</sup> Two fetuses in late stages of degeneration.
D40, gestational day 40; D48, gestational day 48; HB, heartbeat.

Red fluorescence was observed throughout all tissues in the transgenic fetuses (F12 and F13) compared with OFF4 WT (Fig. S9). In healthy mCherry fetuses, the fetal tissue had a purple/red hue, visible to the naked eye. Length and weight of healthy transgenic and WT fetuses were similar (Table S8). Likewise, the area of metanephros (Fig. S10a), gonads (Fig. S10b), adrenals (Fig. S10c), and mesonephros (Fig. S10d) in OFF4 *mCherry54* and OFF4 WT fetuses were comparable. Histological examination of kidneys from healthy transgenic fetuses revealed normal renal architecture, with comparable numbers of stromal cells and the presence of both glomeruli and renal tubules (Fig. S11).

### Kidney complementation

#### Fluorescence-based chimaerism

Chimaeric embryos and fetuses were generated from combining two different TKO host cells (indel vs large deletion, *CGS4^hypo^* and *CGS20^null^*, respectively) with two different mCherry donors (high- vs low-copy number, *mCherry38* vs *mCherry54*, respectively), resulting in three different types of aggregation chimaeras (*CGS4^hypo^*↔*mCherry38*, n=5; *CGS4^hypo^*↔*mCherry54*, *n*=3; and *CGS20^null^*↔*mCherry54*, *n*=3; Table S9). No *CGS20^null^*↔*mCherry38* combinations were tested once it transpired that *mCherry38* derivates were not viable to D48. Over 90% of all aggregation morulae developed into blastocysts. Based on the mCherry fluorescence signal, blastocysts were categorized into three groups: high (likely non-chimaeric, *mCherry* donor embryo only), mixed (chimaeric), or no signal (non-chimaeric, TKO host embryo only). Most blastocysts contained mixed mCherry signal across all strain combinations (*CGS4^hypo^*↔*mCherry38* 79/96=82%, *CGS4^hypo^*↔*mCherry54* 57/65=88%, and *CGS20^null^*↔*mCherry54* 59/59=100%).

Putative chimaeric blastocysts from the three aggregation combinations were transferred into recipients (Table 3, Fig. 4a). All three groups established pregnancies and produced fetuses with a heartbeat with similar efficiency. On D48, fetuses were collected when either a fetal heartbeat was detected or other pregnancy parameters (e.g. fluid levels, fetus and placentome size) were within normal limits. Six fetuses were recovered from *CGS4^hypo^*↔*mCherry38* chimaeras, five were healthy (5/60=8%) and one had arrested around D30 based on CRL. 14 fetuses were collected from *CGS4^hypo^*↔*mCherry54* chimaeras, eight were healthy (8/46=17%) and six in varying stages of degeneration, including one showing early symptoms of fetal hydrops. Five fetuses were collected from *CGS20^null^*↔*mCherry54* chimaeras, two were healthy (2/47=4%) and three were in early or late stages of degeneration. All three combinations produced chimaeric fetuses with donor-derived, mCherry-fluorescent kidneys that appeared anatomically and histologically normal (Fig. 4b). The size of the female donor-derived kidneys varied widely between complementation groups, individuals within each group and even kidneys within each individual (Fig. 4c). For both *CGS4^hypo^*↔*mCherry54* and *CGS20^null^*↔*mCherry54*, chimaeric fetuses with higher mCherry signal tended to be closer to the WT female size. Kidneys with lower fluorescent signal from *CGS4^hypo^*↔*mCherry54* and *CGS20^null^*↔*mCherry54* chimaeras (F36 and F29) were about half the size of female WT controls and ∼70% smaller than male WT. In contrast to *CGS4^hypo^*↔*mCherry54* and *CGS20^null^*↔*mCherry54* chimaeras, all of the *CGS4^hypo^*↔*mCherry38* fetuses had kidneys smaller than the 95% CI of WT females (>60% smaller than male WT, >30% smaller than female WT). Regardless of the complementation group, chimaeric kidneys fell within the range of either exclusively donor- or host-derived organ sizes, but none of them reached the average size of WT male control kidneys. For *CGS4^hypo^*↔*mCherry38*, five healthy fetuses were dissected (Fig. S12). One fetus (F06) was highly red fluorescent in all tissues, three fetuses (F04, F09, F10) showed low levels of red signal that was restricted to the meta- and mesonephroi, while no fluorescence was observed in the F05. For *CGS4^hypo^*↔*mCherry54*, 10 healthy fetuses were dissected (Fig. S13). Four fetuses (F32, F34, F37, and F39) exhibited strong red fluorescence across all tissues examined. A further four fetuses (F31, F33, F35, and F36) showed lower levels of fluorescence throughout all tissues, with the reduction being most apparent in the isolated kidneys. One fetus (F23) displayed fluorescence exclusively in the metanephros and mesonephros, whereas no detectable fluorescence was observed in F22. For *CGS20^null^*↔*mCherry54*, four fetuses were dissected (Fig. S14). All of them (F25, F26, F28, F29) were red fluorescent in all tissues, including the isolated kidneys. Health status, CRL, fetal weight and organ area (kidney, gonads) for all complementation groups are summarized in Table S10. To determine chimaerism based on mCherry, we used the average red fluorescence (normalized on *mCherry54*) of healthy non-transgenic fetuses (derived from *CGS4^hypo^*) as a cut-off; anything above this background value was considered positive, either chimaeric or donor-only (Table S11).

**Fig. 4.**
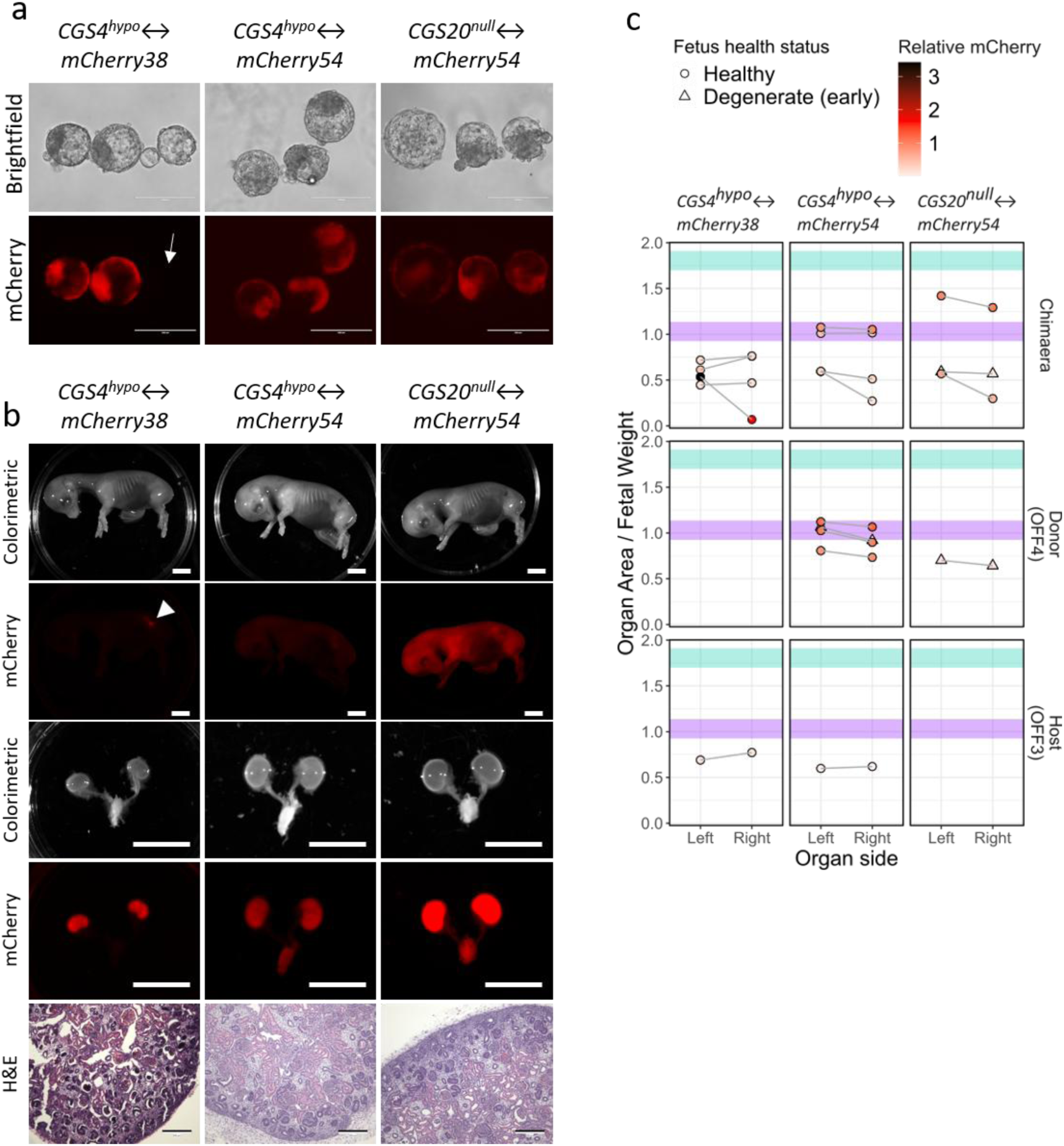
Kidney rescue in chimaeric embryos and fetuses. (a) Brightfield and fluorescent signal in cloned aggregation blastocysts derived from *CGS* host (*4, 20*) and *mCherry* (*38, 54*) donor cells. Fluorescent images were captured at 40% (*mCherry38*) and 60% (*mCherry54*) light intensity. Arrow indicates putative non-chimaeric embryo with no red fluorescence. Scale bar = 200 µm. (b) Representative healthy D48 fetuses derived from cloned aggregation blastocysts as under (a) *CGS4^hypo^*↔*mCherry38* fetus (F10), *CGS4^hypo^*↔*mCherry54* (F31, *CGS20^null^*↔*mCherry54* (F28). Colorimetric images show whole fetus (1^st^, 2^nd^ rows) and isolated metanephros (3^rd^, 4^th^ rows) captured by ChemiDoc (0.2 s exposure on DyLight550 channel). Arrowhead indicates putative donor-derived kidney in chimaeric fetus. scale bar = 10 mm. Histological hematoxylin/eosin (H&E)-stained sections (5^th^ row, scale bar = 200 µm) (c) Normalized metanephros area from collected healthy fetuses (top) *CGS4^hypo^*↔*mCherry38*, (middle) *CGS4^hypo^*↔*mCherry54*, and (bottom) *CGS20^null^*↔*mCherry54*. mCherry fluorescence level indicated by red coloring of circles (relative to *mCherry54*). 95% CI for WT female (lavender) and male (aqua). Paired organs from a single fetus are connected with a grey line.

**Table 3.** In vivo survival of chimaeric cloned embryos.

| Donor cells | Recipient ewes | Embryos transferred | Scan 1 <D40 |  | D48 fetuses |  |
| --- | --- | --- | --- | --- | --- | --- |
|  |  |  | Pregnant (%)<sup>a</sup> | HB (%)<sup>a</sup> | Collected healthy (%)<sup>a</sup> | Collected aborting (%)<sup>a</sup> |
| *CGS4<sup>hypo</sup>↔mCherry38* | 20 | 60 | 15 (25) | 10 (17) | 5 (8) | 1 (2)<sup>b</sup> |
| *CGS4<sup>hypo</sup>↔mCherry54* | 15 | 46 | 12 (26) | 10 (22) | 8 (17) | 6 (13)<sup>c</sup> |
| *CGS20<sup>null</sup>↔mCherry54* | 15 | 47 | 14 (30) | 7 (15) | 2 (4) | 3 (6)<sup>d</sup> |
| **Total** | **50** | **153** | **41 (27)** | **27 (18)** | **15 (10)** | **10 (7)** |

<sup>a</sup> Normalized on blastocysts transferred.
<sup>b</sup> One fetus arrested approximately D30.
<sup>c</sup> One fetus showing signs of fetal hydrops, one in early stages of degeneration, and four in late stages of degeneration.
<sup>d</sup> Two fetuses in early stages of degeneration and one in late stages of degeneration.
D40, gestational day 40; D48, gestational day 48; HB, heartbeat.

#### PCR-based chimaerism

To validate the chimaerism estimates based on mCherry fluorescence, we used qualitative endpoint PCR with donor- and host-specific primers and quantitative ddPCR for *mCherry*. Sensitivity of the assay was determined by combining *CGS4^hypo^* and *mCherry54* DNA to simulate chimaeric contributions from 0.1%–50%, followed by endpoint PCR with female and male-specific primers (Fig. S15ai, ii). Primers designed to detect the female strains were slightly more sensitive than male-specific primers, detecting 0.1% and 0.25% chimaerism, respectively. Fetuses derived from *CGS4^hypo^*↔*mCherry38*, *CGS4^hypo^*↔*mCherry54*, and *CGS20^null^*↔*mCherry54* aggregation were analyzed for presence of the Y chromosome (of the male host) and the -1 bp *CMAH* edits (of the female donor) by using this endpoint PCR assay. For *CGS4^hypo^*↔*mCherry38*, genomic DNA from heart and kidney was only positive for both donor- and host-specific primers in one fetus (F06; Fig. S15b), whereas several chimaeric fetuses were identified from *CGS4^hypo^*↔*mCherry54* (F31, F32, F35, F36; Fig. S15c) and *CGS20^null^*↔*mCherry54* (F26, F28, F29; Fig. S15d).

Using ddPCR, F06, the only chimaeric *CGS4^hypo^*↔*mCherry38* fetus from the endpoint PCR, contained varying *mCherry* contributions throughout heart, kidney, brain, and liver (Fig. S16a). Brain tissue carried *mCherry* copy numbers close to the parental *mCherry38* donor cells, followed by liver, kidney, and heart with lower, but detectable levels of *mCherry* contribution. Of the four healthy *CGS4^hypo^*↔*mCherry54* putative chimaeras, F32 had high levels of *mCherry* contribution to all lineages, while F31, F35, and F36 had lower levels of *mCherry* (Fig. S16b). Another three fetuses (F34, F37, F39) that were not chimaeric in endpoint PCR showed similar signals as the positive control (*mCherry54*) across all four tissues by ddPCR indicating they were solely donor-derived. The three *CGS4^hypo^*↔*mCherry54* fetuses that were negative for female DNA (F22, F23, F24) were also negative for *mCherry* contribution. No *mCherry* DNA was detected in the fetus with fluorescence seen in the kidneys (F23). All *CGS20^null^*↔*mCherry54* putative chimaeras were confirmed using ddPCR across the four tissues (Fig. 16c). The four *CGS4^hypo^*↔*mCherry38* and *CGS4^hypo^*↔*mCherry54* fetuses with low level fluorescence detected in the kidneys, but little to absent *mCherry* DNA were categorized as kidney-specific chimaeras for further analysis (F04, F09, F10, and F23).

Considering all molecular assays, the average chimaerization efficiency was 54%, with no significant differences across the three complementation groups (Table 4). Generally, the fluorescence- and different PCR-based assays were in good agreement (Table S12). When the mCherry fluorescence was above background levels, we relied on the combined PCR assays to discriminate between donor-only derived and chimaeric fetuses.

**Table 4.** In vivo chimaerization efficiency in cloned fetuses.

| Strains | Total fetuses | Chimaeric (%) | Non-chimaeric (%) |
| --- | --- | --- | --- |
| *CGS4<sup>hypo</sup>↔mCherry38* | 5 | 4 (80) | 1 (20) |
| *CGS4<sup>hypo</sup>↔mCherry54* | 14 | 6 (43) | 8 (57) |
| *CGS20<sup>null</sup>↔mCherry54* | 5 | 3 (60) | 2 (40) |
| **Total** | **24** | **13 (54)** | **11 (46)** |

Efficiency normalized on number of total fetuses.

#### Organ analysis

Last, we compared the fluorescence level and size of fetal gonads (Fig. S17a), adrenals (Fig. S17b), and mesonephros (Fig. S17c) in putative chimaeras with WT male host (OFF3) and female donor (OFF4) fetuses.

In *CGS4^hypo^*↔*mCherry54* chimaeras, the four fetuses with the highest mCherry signal (female-only F34, F37, F39, and chimaera F32) had gonads within 10% of the mean WT female size. Gonads in the two chimaeras with lower fluorescence were larger, within the 95% CI for WT males but 25%–57% larger than females. This was similar to the male-only fetuses that were also within 30% of male WT size. Similarly, in *CGS20^null^*↔*mCherry54* fetuses, the chimaera with the highest mCherry signal had gonads close to the expected size of female controls, while chimaeras with lower mCherry signal had larger gonads, within the expected size of males. As for kidney, gonad sizes in *CGS4^hypo^*↔*mCherry38* fetuses were less correlated with mCherry signal, with all five fetuses being close to the 95% CI for male gonads. While left and right gonads differed in size, there was no clear trend for one side to be consistently larger.

Adrenal sizes in both *CGS4^hypo^*↔*mCherry54* and *CGS20^null^*↔*mCherry54* groups showed a similar pattern to gonads, with higher mCherry signal correlating with smaller, “female-like” organ sizes and lower mCherry signal fetuses being closer to the 95% CI for male adrenals. Mesonephros size did not follow the trend for the other three organs with highly fluorescent organs being either similar or larger than female WT controls.

Last, metanephric kidney morphology was qualitatively evaluated on stained sections from each complementation group (Fig. S18). Except for F06, which was cut deeper in the stroma due to different tissue processing, the structure of all chimaeric kidneys was similar to that of WT controls, showing comparable populations of stromal cells, glomeruli, and tubules.

## Discussion

### *SALL1* KO phenotype

Depending on the edit, *SALL1^−/-^* fetuses either produced a variable hypomorph (*CGS4^hypo^*) or lacked kidneys altogether (*CGS20^null^*). *CGS4^hypo^* phenotypes ranged from unilateral agenesis or severe bilateral hypoplasia (62%–92% smaller than WT male) to mild bilateral hypoplasia (20%–40% smaller than WT male), matching the size of female host kidneys. Morphology and histology of hypoplastic kidneys appeared normal in *CGS4^hypo^* fetuses, which contrasts with pig *SALL1*^−/-^ hypoplastic kidneys that displayed disorganized metanephric structure, fewer nephrons, and necrosis at a similar developmental time point (D48 vs D40, in sheep vs pig, respectively) (Watanabe et al., 2019). One *CGS4^hypo^* fetus was missed at D48 scanning and collected on D77. In this fetus, the left kidney was lower and medially positioned compared to the right kidney, similar to the abnormal kidney anatomy observed in a *SIX1*^−/-^ pig fetus with mild bilateral hypoplasia (J. Wang et al., 2019). The described fetal phenotypes suggest a partial loss-of-function for SALL1 since no bilateral lack of kidney was observed. The single gRNA cut occurred 5’ of ZF5, leaving the first four C2HC-type ZFs intact. A similar approach had been taken previously with a single guide targeting 5’ of ZF3 in pig (Kim et al., 2016), but no phenotype was reported for this mutation. In addition to ZFC1 and most of ZFC2, this edit also left three other functional SALL1 domains intact, namely: i) a conserved 12 amino acid motif at the N-terminus that interacts with the nucleosome remodeling and histone deacetylase (NuRD) complex (Lauberth & Rauchman, 2006), ii) a C2HC-type ZF domain (Álvarez et al., 2021), and iii) a glutamine-rich (polyQ) domain necessary to mediate spalt protein interactions and subcellular localization (Sweetman et al., 2003). Collectively, this leaves several functional domains in place to mediate both residual transcriptional repression and heterochromatin binding (Netzer et al., 2006), consistent with the observed phenotype.

To achieve full kidney agenesis, new Cas9 gRNAs were designed to delete the entire exon. This approach had produced anephric mice (Nishinakamura et al., 2001) and pigs (Watanabe et al., 2019). This second round of editing was performed on rejuvenated *CGS4^hypo^* cells, which had produced a viable D77 fetus with kidneys. It involved two gRNAs to delete all C2H2-type ZFs, as well as the C2HC- type ZF and polyQ domain through a large frameshift mutation (Cong et al., 2013). This mutant only retained the N-terminal NuRD region, which may mediate residual transcriptional repression, but no longer localizes to the heterochromatin (Netzer et al., 2006). This mutant mimics the original editing in anephric mice (Nishinakamura et al., 2001). Only one fetus developed to D48, which was likely an SCT cloning artefact, rather than being caused by the deletion because *SALL^−/-^*pigs (Watanabe et al., 2019) and rodents (Goto et al., 2019; Nishinakamura et al., 2001) survived to D40 and term, respectively. Given the small numbers of embryos transferred, no conclusions can be drawn as to whether the mutation in *CGS20^null^* affected pregnancy rate. The single fetus collected on D48 lacked both kidneys, matching the phenotype of rodent and pig *SALL1^−/-^*. More *CGS20^null^* fetuses are needed to confirm the KO phenotype and determine any possible phenotypic variability.

Assessing pleiotropic effects of *SALL1^null^* is important for determining its viability for use in the organ niche model. Humans with heterozygous *SALL1* mutations develop Townes-Brocks syndrome, which yields malformations in renal, anal, limb, and ear development (Kohlhase et al., 1998). However, no abnormal development was observed in these systems, or other areas of normal *SALL1* expression, in heterozygous or homozygous mice mutants (Nishinakamura et al., 2001). It is hypothesized that overlapping expression of other *SALL* family genes (*SALL2* and *SALL3*) may compensate development in other organs where *SALL1* is expressed. Even though the *CGS20^null^*fetus was in the early stages of degeneration, no other developmental abnormalities were observed, indicating that sheep may be a viable organ niche model for this mutation.

In contrast to rodent and pig studies, both our mutant *SALL1* lines were generated on a background of identical *CG^null^*xenocompatible mutations. The frameshift mutations in *CMAH* had been characterized previously to prevent the formation of Neu5Gc. One edit in *GGTA1* was a frameshift, while the other allele deleted 101 amino acids within the functional domain, which likely abolished function of the protein. We observed no renal or other organ abnormalities in live *CG^null^* sheep (Appleby et al., 2026) and there are no known interactions between SALL1 and the surface antigens generated by CMAH and GGTA1. Thus, it is unlikely that removing the xenoantigens would have affected the *SALL1^null^* phenotype.

### Producing fetal chimaeras

We previously established aggregation of cloned host and donor morulae as an optimal strategy for producing sheep chimaeras (McLean et al., 2021, 2025). Here we generated three chimaeric combinations by aggregating hypomorph host *CGS4^hypo^*with donor mCherry strains (*mCherry38* and *mCherry54*, respectively), and full KO host *CGS2^null^* with *mCherry54*. Chimaerism was determined by three independent assays — mCherry fluorescence, *mCherry* ddPCR, and donor-specific endpoint PCR — which revealed 60% of fetuses as chimaeric and 40% as donor-only. The overall level of fetal chimaerism and kidney complementation was 54% (13/24) with no significant differences across the three chimaera combinations. This was the same efficiency as in our study of neonatal chimaerism and germline complementation in sheep using a wild-type low-copy *mCherry* donor strain from the same parental line (7/13=54%) (McLean et al., 2025). Chimaerism was higher than the 11%–36% previously observed in pigs (Ji et al., 2017; Nakano et al., 2013), but it was not reported if non-chimaeric porcine blastocysts were screened out by absence of fluorescence before transfer, potentially lowering chimaerism efficiency.

High-copy donor *mCherry38* was embryonic-lethal, likely due to random integration into vital genomic sites and high mCherry expression impairing in vivo development (Shemiakina et al., 2012).

Consequently, no kidney rescue was observed in *CGS4^hypo^*↔*mCherry38* chimaeras. A similar observation was made with pig chimaeras where the anephric phenotype was not rescued due to low donor cell contribution (Matsunari et al., 2020). By contrast, both *mCherry54* chimaeric combinations had normal organ sizes and morphology, matching OFF4 WT controls. Therefore, further discussion focuses on *CGS20^null^*↔*mCherry54* chimaeras.

### Rescued kidney phenotype

In one chimaera (F28), kidneys were approximately normal size, falling between the 95% CI for male and female WT fetuses. Kidneys in the second healthy fetus (F29) were smaller, indicating partial rescue. The remaining fetuses had stalled development and were degenerating, which likely explains their bilateral kidney hypoplasia. Gonad size followed a similar pattern to kidneys in the healthy fetuses. The highly fluorescent chimaera contained female-sized gonads, confirming this fetus was mostly derived from *mCherry54* donor cells, with low contribution of male *CGS20^null^* hosts. The chimaeric fetus (F29) with lower mCherry contribution contained male-sized gonads, in line with the male KO host being the dominant cell population. Degenerating chimaeric and female-only fetuses matched expected size for WT females.

Female sized kidneys could mean that either the kidneys were fully rescued by the female donor cells to correct female size, or the kidneys were displaying mild hypoplasia being ∼40% smaller than normal kidneys for the male host. In the second situation, hypoplasia may have resulted from compromised development of cells expressing mCherry. Sexual dimorphism was unexpected, as there are no reports in the literature indicating that this occurs. An alternative to size differences based on sex was that the male and female parental cell lines were different breeds. Both cell lines were composite breeds, with predominantly Perendale contribution in the male OFF3 and Coopworth in the female OFF4. Both dominant breeds are average sized sheep, suggesting breed was unlikely to affect organ size. Based on mouse↔rat chimaeras it was expected that resulting organs would match the size of the host (Kobayashi et al., 2010; Yamaguchi et al., 2017). Avoiding intersex chimaeras in future experiments would aid analysis of whether rescued organs were normal or exhibiting mild hypoplasia, but complicate subsequent breeding.

The cloning process itself may have also compromised kidney rescue by introducing epigenetic errors. Cloned pig donors failed to rescue the anephric phenotype of cloned *SALL1* KO hosts but using IVF-derived embryos led to 1 chimeric fetus of 12 with morphologically and histologically normal kidneys (Matsunari et al., 2020). Confirmation and improvement in the efficiency of complete kidney rescue, e.g. using IVF donor embryos to improve donor cell survival, is required before attempting to take chimaeras further into gestation, and eventually to term.

### Producing xenocompatible organs in livestock

Producing human organs in livestock could bypass issues of incompatible functioning of livestock organs. The first step towards this goal is to generate viable intraspecies livestock chimaeras, in which the organ of interest is comprised entirely of the donor cells. This would then be followed by closely related interspecies chimaeras, and finally human↔livestock chimaeras growing the right-sized, functional organ. To produce heterologous kidneys in intraspecific chimaeras, we used female donor and male host strains, so resulting intersex chimaeras would be phenotypically male. If chimaeras were to survive to term with fully rescued kidneys, the resulting animals would generate sperm solely from the male *SALL1*^−/-^ host strain. An alternative would be to use *DAZL* or *NANOS2* KO donor cells (McLean, Appleby, Wei, et al., 2021; McLean et al., 2025; Park et al., 2017), which could fill the empty kidney niche without contributing to the male germline.

Live chimaeras with complemented kidneys would be an important milestone as rodent chimaeras died shortly after birth, associated with abnormal development of the olfactory bulb (Nishinakamura et al., 2001). In pigs, homozygous *SALL1* KO is prenatally lethal sometime after D40 (Watanabe et al., 2019). If this phenotype was carried into other large animal models, this would complicate future work using *SALL1* for kidney complementation. Conditional KOs may be required to ensure loss of *SALL1* was restricted to the kidney to alleviate pleiotropic effects.

Another limitation to producing heterologous *SALL1* WT kidneys is that structures derived from the ureteric bud (collecting ducts, ureter, renal pelvis), and microvasculature are still chimaeric. To limit contribution of host cells within the kidney, further genetic modifications would be required. To remove host cells from ureteric bud structures, loss-of-function mutations could be introduced to both FGF and GDNF signaling components, which are both involved in initiating metanephros formation from the ureteric bud. Candidate target genes to prevent vascularization could be added alongside other organogenesis gene KOs (Hamanaka et al., 2018; Matsunari et al., 2020).

Blastocyst complementation has generated a range of solid organs (liver, lung, kidney, pancreas, heart, thyroid, thymus, and parathyroids), but only a subset of these were generated in livestock (kidney, liver, pancreas), namely pig (Bigliardi et al., 2025). Apart from rat–mouse chimaeras, there is only one interspecies model that used human donor cells in a pig host (J. Wang et al., 2023). Wang et al. complemented *SIX1* and *SALL*1 DKOs with DsRed-labelled human iPSCs and recovered embryos on D25 or D28 (J. Wang et al., 2023). While no fully developed kidneys were obtained, the mesonephros in the chimeric embryos was histologically similar. Furthermore, over 50% of the mesonephric cells showed DsRed expression in both mesonephric tubules and mesenchymal cells, as well as expressing several kidney developmental markers (*SALL1, SIX1, PAX2,* and *WT1*). Despite this progress, major hurdles for successful interspecies chimaerism remain, including donor-host (i) cell competition, (ii) ligand and receptor incompatibility, (iii) asynchrony in developmental speed, and (iv) mismatch in developmental stages at the time of embryo complementation (Bigliardi et al., 2025).

In addition to these biological barriers, repurposing animals as disposable, commodified “organ factories” raises ethical concerns about animal rights and welfare, as well as around the potential of human cells to contribute to cognition, gametogenesis, and appearance (Behnam Manesh et al., 2014). The latter may be alleviated by additional genetic modifications, but public perception of these risks will need addressing. Additional concerns around the potential for inequality based on costly organ production would also need to be addressed. Last, the use of pigs in particular raises cultural and religious implications. Many Islamic cultures prohibit transplantation of porcine organs to replace defective human ones (Mohd Zailani et al., 2023), even though they may be permissive if it was a lifesaving measure (Paris et al., 2018). A similar argument would apply to Jewish patients, even though they can more freely use porcine products as lifesaving organ transplants (Rosner, 1999). Using sheep as an alternative donor species might mitigate some of these issues.

## Acknowledgments

We thank the Ruakura farm staff: Aaron Malthus and Tim Hale took care of animal husbandry, while Elyssa Barnaby and Ali Cullum provided veterinarian care. Ovary collection was performed by Murray Brown, Angela Brennan, Tony Kerema, Katherine Moors, and Sue Odom. Cas9 plasmids were kindly provided by the Zhang lab through Addgene and the CAGGS-mCherry and SB100 plasmids were donated by Wilfried Kues. Stefan Wagner designed and provided the CSN2 probe. The Davidson lab at the University of Auckland: Dr George Chang, Dr Jennifer Hollywood, Dr Zhenzhen Peng, and Dr Veronika Sander provided technical help and guidance and Aparajita Chatterjee and Tayla Perreau aided with histology. Thanks to Brigid Brophy and Dr Blaise Forrester-Gauntlett for helpful comments on the manuscript. This work was funded by a PhD fellowship from the University of Auckland to S.J.A., with further support by the Todd Foundation and Maurice & Phyllis Paykel Trust.

## Author Contributions

S.J.A. and B.O. conceived and designed the experiments. S.J.A. carried out the experiments, processed and analyzed the data, and produced the figures. L.M.F. aided in fetal dissections, sheep cloning, and project administration. S.D. took care of embryo transfers and animal husbandry. J.W., F.M., and D.N.W. contributed to sheep cloning and cryopreservation. P.T. aided in ddPCR analysis. A.J.D. provided supervision and histology resources. S.J.A. and B.O. wrote the manuscript. All authors read, provided comments, and approved the final manuscript.

