## Supplemental Figures and Tables for "Morula complementation restores fetal kidneys in xenocompatible *SALL1* null sheep"

*corresponding author

### Supplementary Tables

**Table S1.** Sequences of gRNAs, oligos, and primers.

| Name | Target | Direction | Sequence | Amplicon size (bp) |
|---|---|---|---|---|
| **gRNA1** | SALL1 | f | caccgTAGCTCGATGGACGCTGTAG | |
| **gRNA1** | SALL1 | r | aaacCTACAGCGTCCATCGAGCTAc | |
| **gRNA2** | SALL1 | f | caccgTGAGCGGGGGCATAGCTCGA | |
| **gRNA2** | SALL1 | r | aaacTCGAGCTATGCCCCCGCTCAc | |
| **gRNA3** | SALL1 | f | caccgAGCTCGATGGACGCTGTAGT | |
| **gRNA3** | SALL1 | r | aaacACTACAGCGTCCATCGAGCTc | |
| **gRNA4** | SALL1 | f | caccgCAACCGCACCACTAAGAGCA | |
| **gRNA4** | SALL1 | r | aaacTGCTCTTAGTGGTGCGGTTGc | |
| **gRNA5** | SALL1 | f | caccGAGTTCTCTCGTGAATCTGC | |
| **gRNA5** | SALL1 | r | aaacGCAGATTCACGAGAGAACTC | |
| **BJO289** | SALL1 | f | CTTTGGGGGACTCTTGGACTC | 421 |
| **BJO290** | SALL1 | r | CTCATCAAAGGAGCCCGTGTC | |
| **BJO409** | SALL1 gRNA4 FWD | f | TGCACCAATGTCTGCAGTTC | 729 |
| **BJO410** | SALL1 gRNA4 REV | r | AGGTCCCCGAGTTGAGGTAG | |
| **BJO413** | SALL1 gRNA5 FWD | f | TGACACATCAGATGCGGGAT | 482 |
| **BJO414** | SALL1 gRNA5 REV | r | ACTCTGGGAGCGTGCTTTAT | |
| **BJO380** | SALL1 gRNA2 off target 1 | f | CCTTCTGGCCCTAAAATCTTCAGT | 792 |
| **BJO381** | SALL1 gRNA2 off target 1 | r | CAGGCAATCTGGCTCCTAGTC | |
| **BJO382** | SALL1 gRNA2 off target 2 | f | AGACTTTGCGATTCTCCCCG | 852 |
| **BJO383** | SALL1 gRNA2 off target 2 | r | AGAGAGGAAGGGGCTCTCTG | |
| **BJO390** | SALL1 gRNA2 off target 3 | f | AGGGTGTGGATGCTCTGTTG | 693 |
| **BJO391** | SALL1 gRNA2 off target 3 | r | TTAGAAAGCCACTGCCCCAG | |
| **BJO418** | SALL1 gRNA4 off target 1 | f | CCCAACTCCACGCTTGATCT | 472 |
| **BJO419** | SALL1 gRNA4 off target 1 | r | GGAAGGGTCTTCAGTTCGCA | |
| **BJO420** | SALL1 gRNA4 off target 2 | f | TAGCTTTCCTTTGGGTGCATGT | 573 |
| **BJO421** | SALL1 gRNA4 off target 2 | r | AAGGAACTGAATCAGAAGAGGCT | |
| **BJO422** | SALL1 gRNA4 off target 3 | f | AAGCACCTTGAAATGGCCCC | 535 |
| **BJO423** | SALL1 gRNA4 off target 3 | r | GGGGTCGGGACATTCTACTC | |
| **BJO424** | SALL1 gRNA5 off target 1 | f | CATTGGGCTTTTCTTGGCATTT | 604 |
| **BJO425** | SALL1 gRNA5 off target 1 | r | AGCAACCTTCCTGCTAAGGC | |
| **BJO426** | SALL1 gRNA5 off target 2 | f | GCCCCTTCATGCCTGATTTG | 643 |
| **BJO427** | SALL1 gRNA5 off target 2 | r | ATGTGGCTGTAGAGAGGGCA | |
| **BJO428** | SALL1 gRNA5 off target 3 | f | CTTTCCACAAGCCCTCGTCT | 596 |
| **BJO429** | SALL1 gRNA5 off target 3 | r | GCTCCTCGGAGCTCTGAATC | |
| **BJO392** | mCherry CNV Fam probe | | CAGTACGAACGCGCCGAGGG | 133 |
| **BJO393** | mCherry CNV assay | f | AACATCAAGTTGGACATCAC | |
| **BJO394** | mCherry CNV assay | r | TTGAGCTCGAGATCTGAG | |
| **BJO468** | Female mCherry CMAH | f | CATTCAGTGGCTATTTCCAGCC | 471 |
| **BJO469** | Female mCherry CMAH | r | AGGACCCGTTAACAAGGGTCAAA | |
| **DDX3Y-F** | Male DDX3Y | f | GGACGTGTAGGAAACCTTGG | 685 |
| **DDX3Y-R** | Male DDX3Y | r | GCCAGAACTGCTACTTTGTCG | |
| **GL465** | Cas9 plasmid CBh | r | GGGCAGTTTACCGTAAATAC | 328 |
| **GL1358** | Cas9 insertion | f | CAGAGCTTCATCGAGCGGAT | 466 |
| **GL1359** | Cas9 insertion | r | CGAACAGGTGGGCATAGGTT | |
| **GL1041** | Plasmid sequencing | | ACTATCATATGCTTACCGTAAC | |
| **M13_f** | Plasmid sequencing (Massey) | f | CCCAGTCACGACGTTGTAAAACG | |

Plasmid overhangs indicated with lowercase letters. Underlined primers used for sequencing.

**Table S2.** *SALL1* gRNA sequences and efficiency scores from CRISPOR.

| gRNA number | PAM | Specificity score | Predicted efficiency | Out-of-frame score | Predicted off-target sites |
|---|---|---|---|---|---|
| gRNA1 | TGG | 95 | 50 | 70 | 12 |
| gRNA2 | TGG | 96 | 60 | 75 | 41 |
| gRNA3 | GGG | 96 | 52 | 70 | 29 |
| gRNA4 | AGG | 86 | 61 | 63 | 78 |
| gRNA5 | AGG | 84 | 48 | 78 | 75 |

**Table S3.** Mutation profiles of triple knockout (TKO) clonal strains.

| Cell line | Strain number[a] | *CMAH* Edit 1 | *CMAH* Edit 2 | *GGTA1* Edit 1 | *GGTA1* Edit 2 | *SALL1* Edit 1 | *SALL1* Edit 2 | Frameshift all genes | Cas9 present | Off target sequence[b] | SCT |
|---|---|---|---|---|---|---|---|---|---|---|---|
| OFF3 | 4 | -34 bp | -44 bp | +1 bp | -303 bp | +1 bp | -11 bp | Yes | No | WT | Yes |
| | 5 | +1 bp | -396 bp | +1 bp | +1 bp | No PCR band[c] | | Yes | No | WT[d] | |
| | 6 | -4 bp | -15 bp | +1 bp | -1 bp | -125 bp | -125 bp | | | | |
| | 9 | -3 bp | -29 bp | +1 bp | -1 bp | +1 bp | -14 bp | | | | |
| | 10 | -12 bp | -13 bp | -1 bp | -3 bp | +1 bp | +1 bp | | | | |
| | 11 | -1 bp | -3 bp | -1 bp | -6 bp | +1 bp | +1 bp | | | | |
| | 12 | -1 bp | -6 bp | +1 bp | +1 bp | +1 bp | +1 bp | | | | |
| | 13 | -3 bp | -3 bp | +1 bp | -6 bp | -14 bp | -14 bp | | | | |
| | 14 | -7 bp | -7 bp | +1 bp | -1 bp | -12 bp | -12 bp | | | | |
| | 15 | -3 bp | -11 bp | -1 bp | -4 bp | -7 bp | -7 bp | | | | |
| | 16 | -5 bp | -18 bp | +1 bp | -2 bp | -44 bp | -44 bp | | | | |
| | 17 | -3 bp | -13 bp | -1 bp | -19 bp | +1 bp | +1 bp | | | | |
| | 19 | +1 bp | +1 bp | +1 bp | +1 bp | +1 bp | -1 bp | Yes | No | WT | |
| | 21 | -26 bp | -26 bp | +1 bp | -1 bp | +1 bp | +1 bp | Yes | No | WT | |
| OFF3 *CGS4*$^{hypo}$ | 2 | -34 bp | -44 bp | +1 bp | -303 bp | -2745 bp | -3308 bp | | | | |
| | 3 | -34 bp | -44 bp | +1 bp | -303 bp | -3309 bp | -3309 bp | | | | |
| | 14 | -34 bp | -44 bp | +1 bp | -303 bp | -2745 bp | -3308 bp | | | | |
| | 20 | -34 bp | -44 bp | +1 bp | -303 bp | -3274 bp | -3274 bp | | No | WT | Yes |

[a] Strains 1, 2, 3, 7, 8, 18, and 20 were mixed clones and are not shown.
[b] Sequences in strain matched wild-type (WT) for that cell line.
[c] No PCR band amplified for *SALL1.*
[d] Analysis performed only for *CMAH* and *GGTA1*.
Cas9 plasmid insertion and off-target screening reported for strains tested.
*CGS4*$^{hypo}$ = *CMAH*$^{-/-}$ *GGAT1*$^{-/-}$ *SALL1*$^{-/-}$ strain #4; *CMAH* and *GGTA1* edits identical to row 1 (grey text).
SCT, somatic cell transfer.

**Table S4.** Cloned embryo in vitro development from TKO donor cells.

| Donor cells | n[a] | nIVC[b] | >1-cell (%)[c] | Non-aggregate blastocysts | Aggregate blastocysts | B 1-3 (%)[d] | B 1-2 (%)[e] |
|---|---|---|---|---|---|---|---|
| OFF3 *CGS4$^{hypo}$* | 2 | 409 | 312 (76)[B] | 6 | 44 | 50 (16)[B] | 21 (42) |
| OFF3 *CGS20$^{null}$* | 2 | 428 | 385 (90)[A] | 18 | 68 | 86 (22)[C] | 45 (52) |
| OFF4 | 3 | 582 | 427 (73)[B] | 0 | 19 | 19 (4)[A] | 7 (37) |
| **Total** | **7** | **1419** | **1124 (79)** | **24** | **131** | **155 (14)** | **73 (47)** |

[a] Number of somatic cell cloning runs.
[b] Number of embryos placed into in vitro culture (IVC).
[c] Cleavage normalized on nIVC.
[d] Total blastocyst (B 1-3) development normalized on >1-cell embryos.
[e] Proportion of grade 1-2 blastocysts (B 1-2) normalized on total number of blastocysts.
[A,B,C] Values within each column differ with $p$ <0.001.
[B,C] Values within each column differ with $p$ =0.045.

**Table S5.** Summary of health and organ measurements in TKO fetuses.

| Cell line | Edited genes | Strain | Fetus ID | Fetus/ewe | Age | Health status | CRL (mm) | Weight (g) | Left Kidney ($mm^2$) | Right Kidney ($mm^2$) | Left Gonad ($mm^2$) | Right Gonad ($mm^2$) |
|---|---|---|---|---|---|---|---|---|---|---|---|---|
| OFF3 | *CMAH GGTA1 SALL1* | 4 | F01 | single | D48 | Healthy | 86.71 | 13.68 | 1.90 | 9.29 | 5.87 | 6.27 |
| | | | F02 | single | D48 | Healthy | 87.99 | 18.27 | 0.00 | 7.06 | 5.54 | 5.56 |
| | | | F07 | single | D48 | Healthy | 84.39 | 16.39 | 22.70 | 24.30 | 9.98 | 10.17 |
| | | | F08 | twin | D48 | Degenerate (early)[a] | 82.00 | 21.09 | 21.10 | 24.87 | 6.40 | 6.73 |
| | | | F08t | twin | D48 | Degenerate (D30)[b] | 29.93 | 0.73 | | | | |
| | | | F11 | single | D77 | Healthy | 219.00 | 195.31 | 120.00 | 149.81 | 15.25 | 21.02 |
| OFF3 | *CMAH GGTA1 SALL1* | 20 | F21 | single | D48 | Degenerate (early)[a] | 78.51 | 9.83 | 0.00 | 0.00 | 2.85 | 3.15 |
| OFF4 | - | WT | F14 | single | D48 | Healthy | 81.20 | 13.4 | 14.66 | 13.51 | 3.49 | 4.08 |
| | | | F18 | triplet | D48 | Healthy | 84.75 | 15.09 | 20.73 | 18.99 | 3.67 | 3.76 |
| | | | F19 | triplet | D48 | Healthy | 75.70 | 10.16 | 9.09 | 9.01 | 2.35 | 2.74 |
| | | | F20 | triplet | D48 | Healthy | 68.49 | 8.47 | 10.15 | 9.57 | 1.23 | 2.22 |
| OFF3 | - | WT | F15 | single | D48 | Healthy | 80.34 | 11.77 | 20.40 | 22.01 | 5.33 | 4.52 |
| | | | F17 | single | D48 | Healthy | 82.62 | 16.32 | 30.21 | 28.79 | 8.75 | 9.06 |

[a] Fetus in early stages of degeneration.
[b] Fetus arrested at approximately D30.
CRL, crown rump length; WT, wild-type.

**Table S6**. Mutation profiles and Cas9 insertion of *CMAH/GGTA1*-edited OFF4 strains.

| Cell line | Mutation profile | *CMAH* | | *GGTA1* | | | Individual strains isolated | Clonal strain | Frameshift both genes | Cas9 present | Off target sequence[a] | ID |
|---|---|---|---|---|---|---|---|---|---|---|---|---|
| | | Edit 1 | Edit 2 | Edit 1 | Edit 2 | Edit 3 | | | | | | |
| OFF4 | A | -7 bp | -9 bp | -6 bp | -6 bp | | 3 | Yes | No | No | | |
| | B | -3 bp | -3 bp | -3 bp | -6 bp | | 1 | Yes | No | No | | |
| | C | -9 bp | -9 bp | -123 bp | -111 bp | | 2 | Yes | No | No | | |
| | D | +1 bp | -44 bp | 0 bp | -16 bp | | 1 | Yes | No | No | | |
| | E | -27 bp | -27 bp | 0 bp | -24 bp | | 2 | Yes | No | No | | |
| | F | -1 bp | -19 bp | -3 bp | -3 bp | | 2 | Yes | No | No | WT | *mCherry38* |
| | G | -90 bp | -90 bp | -32 bp | -32 bp | | 2 | Yes | No | No | | |
| | H | -7 bp | -20 bp | -2 bp | +1000 bp[b] | | 1 | Yes | ND[d] | No | | |
| | I | 0 bp | 0 bp | 0 bp | +1 bp | | 1 | Yes | No | No | | |
| | J | +1 bp | +1 bp | -12/-13 bp[c] | -37/-38 bp[c] | | 1 | Yes | ND[d] | Yes | WT | |
| | K | -1 bp | -11 bp | 0 bp | -4 bp | | 1 | Yes | No | No | WT | *mCherry54* |
| | M[e] | 0 bp | -27 bp | 0 bp | +1 bp | -24 bp | 1 | No | No | No | | |

[a] Sequences in strain matched wild-type (WT) for that cell line.
[b] Insert was not fully sequenced. Size estimate based on band of PCR product.
[c] Single “A” insertion 5’ cut site on one allele.
[d] Not determined due to inaccuracies in sequencing.
[e] Likely mix of mutation profiles E and I.

**Table S7**. Cloned embryo in vitro development from mCherry donor cells.

| Donor cells | n[a] | nIVC[b] | >1-cell (%)[c] | Non-aggregate blastocysts | Aggregate blastocysts | B 1-3 (%)[d] | B 1-2 (%)[e] | Absolute mCherry fluorescence[f] | Fluorescence relative to *mCherry54* |
|---|---|---|---|---|---|---|---|---|---|
| *mCherry38* | 2 | 370 | 267 (72) | 1 | 31 | 32 (12) | 16 (50) | 2602 | 2.98 |
| *mCherry54* | 4 | 761 | 572 (75) | 11 | 60 | 71 (12) | 38 (54) | 875 | 1.00 |
| **Total** | **6** | **1131** | **839 (74)** | **12** | **91** | **103 (12)** | **54 (52)** | | |

[a] Number of somatic cell cloning runs from *mCherry38* and *mCherry54* donor cells.
[b] Number of embryos placed into in vitro culture (IVC).
[c] Cleavage normalized on nIVC.
[d] Total blastocyst (B 1-3) development normalized on > 1-cell embryos.
[e] Proportion of grade 1-2 blastocysts (B 1-2) normalized on total number of blastocysts.
[f] Fluorescence measured as intensity/$mm^2$. Blastocysts assessed $n$=3.

**Table S8.** Summary of health and organ measurements in mCherry fetuses.

| Cell line | Modified genes | Strain | Fetus ID | Fetus/ewe | Age | Health status | CRL (mm) | Weight (g) |
|---|---|---|---|---|---|---|---|---|
| OFF4 | *CMAH GGTA1 mCherry* | *mCherry54* | F12 | twin | D48 | Healthy | 85.03 | 16.26 |
| | | | F13 | twin | D48 | Healthy | 79.41 | 14.66 |
| | | | F16 A | twin | D48 | Degenerate (late)[a] | 80.56 | 11.91 |
| | | | F16 B | twin | D48 | Degenerate (late)[a] | 69.59 | 7.73 |
| OFF4 | | WT | F14 | single | D48 | Healthy | 81.20 | 13.4 |
| | | | F18 | triplet | D48 | Healthy | 84.75 | 15.09 |
| | | | F19 | triplet | D48 | Healthy | 75.70 | 10.16 |
| | | | F20 | triplet | D48 | Healthy | 68.49 | 8.47 |

[a] Fetus in late stages of degeneration.

CRL, crown rump length; WT, wild-type.

**Table S9**. Cloned embryo in vitro development from aggregation chimaeras.

| Donor cells | n[a] | nIVC[b] | B 1-3 (%)[c] | B 1-2 (%)[d] | Blastocyst mCherry signal (%)[d] | | | |
|---|---|---|---|---|---|---|---|---|
| | | | | | None (host only) | Mixed (chimaeric) | High (donor only) | Not assessed[e] |
| *CGS4*$^{hypo}$↔*mCherry38* | 5 | 93 | 96 (103) | 39 (41)[A] | 11 (11)[C] | 79 (82) | 6 (6) | |
| *CGS4*$^{hypo}$↔*mCherry54* | 3 | 69 | 65 (94) | 30 (46) | 5 (8) | 57 (88) | 0 (0) | 3 (5) |
| *CGS20*$^{null}$↔*mCherry54* | 3 | 58 | 59 (102) | 38 (64)[B] | 0 (0)[D] | 59 (100) | 0 (0) | |
| **Total** | **11** | **220** | **220 (100)** | **107 (49)** | **16 (7)** | **195 (89)** | **6 (3)** | **3 (1)** |

[a] Number of somatic cell cloning runs involving aggregation of male OFF3 *CGS4*$^{hypo}$ or *CGS20*$^{null}$ with female OFF4 *mCherry38* or *mCherry54*.
[b] Number of aggregate embryos placed into in vitro culture (IVC).
[c] Total blastocyst (B 1-3) development normalized on nIVC.
[d] Proportion of blastocysts normalized on total number of blastocysts. Aggregated morulae occasionally gave rise to more than one blastocyst.
[e] mCherry signal not assessed as blastocysts had collapsed.
[A,B] Values within each column differed with p =0.0065
[C,D] Values within each column differed with p =0.0082

**Table S10.** Summary of health and organ measurements in TKO↔*mCherry* aggregation fetuses.

| Cell line | Modified genes | Strain | Fetus ID | Fetus/ewe | Age | Health status[a] | CRL (mm) | Weight (g) |
|---|---|---|---|---|---|---|---|---|
| OFF3 ↔ OFF4 | *CMAH GGTA1 SALL1* ↔ *CMAH GGTA1 mCherry* | *CGS4$^{hypo}$* ↔ *mCherry38* | F03 | single | D48 | Degenerate (D30) | 30.31 | 0.65 |
| | | | F04 | twin | D48 | Healthy | 83.00 | 13.73 |
| | | | F05 | twin | D48 | Healthy | 78.66 | 11.37 |
| | | | F06 | single | D48 | Healthy | 78.31 | 14.24 |
| | | | F09 | single | D48 | Healthy | 86.25 | 16.71 |
| | | | F10 | single | D48 | Healthy | 82.98 | 13.42 |
| OFF3 ↔ OFF4 | *CMAH GGTA1 SALL1* ↔ *CMAH GGTA1 mCherry* | *CGS4$^{hypo}$* ↔ *mCherry54* | F22 | triplet | D48 | Healthy | 88.57 | 12.87 |
| | | | F23 | triplet | D48 | Healthy | 78.81 | 11.72 |
| | | | F24 | triplet | D48 | Degenerate (late) | 66.11 | 5.696 |
| | | | F27 | single | D49 | Degenerate (late) | 78.46 | 13.44 |
| | | | F31 | single | D48 | Healthy | 85.48 | 17.66 |
| | | | F32 | twin | D48 | Healthy | 80.16 | 12.05 |
| | | | F33 | twin | D48 | Degenerate (early) | 80.02 | 14.58 |
| | | | F34 | twin | D48 | Healthy | 89.38 | 16.54 |
| | | | F35 | twin | D48 | Hydrops | 91.78 | 16.2 |
| | | | F36 | twin | D48 | Healthy | 85.00 | 15.06 |
| | | | F37 | twin | D48 | Healthy | 83.19 | 13.96 |
| | | | F38 A | twin | D48 | Degenerate (late) | 70.64 | 9.04 |
| | | | F38 B | twin | D48 | Degenerate (late) | 60.63 | 5.24 |
| | | | F39 | twin | D48 | Healthy | 81.38 | 13.81 |
| OFF3 ↔ OFF4 | *CMAH GGTA1 SALL1* ↔ *CMAH GGTA1 mCherry* | *CGS20$^{null}$* ↔ *mCherry54* | F25 | single | D48 | Degenerate (early) | 89.62 | 19.58 |
| | | | F26 | single | D48 | Degenerate (early) | 93.51 | 19.52 |
| | | | F28 | triplet | D48 | Healthy | 86.38 | 15.62 |
| | | | F29 | triplet | D48 | Healthy | 84.43 | 15.78 |
| | | | F30 | triplet | D48 | Degenerate (late) | 76.74 | 11.89 |

[a] Degenerate fetuses arrested at approximately D30 or in early or late stages of degeneration.
CRL, crown rump length.

**Table S11.** Background red fluorescence in non-chimaeric fetuses.

| Cell line | Modified genes | Strain | n | Health status | mCherry relative to *mCherry54*[a] | | | | | |
|---|---|---|---|---|---|---|---|---|---|---|
| | | | | | Fetus | | | Kidney | | |
| OFF4 | *CMAH GGTA1 mCherry* | *mCherry54* | 2 | Healthy | 1.00 | ± | 0.08 | 1.00 | ± | 0.03 |
| | | | 2 | Degenerate (late) | 0.07 | ± | 0.02 | | | |
| OFF4 | - | WT | 4 | Healthy | 0.01 | ± | 0.00 | 0.02 | ± | 0.00 |
| OFF3 | - | WT | 2 | Healthy | 0.01 | ± | 0.00 | 0.01 | ± | 0.00 |
| OFF3 | *CMAH GGTA1 SALL1* | *$CGS4^{hypo}$* | 3 | Healthy | **0.05** | **±** | **0.01** | **0.08** | **±** | **0.05** |
| | | *$CGS4^{hypo}$* | 1 | Degenerate (early) | 0.04 | | | 0.04 | ± | 0.00 |
| | | *$CGS20^{null}$* | 1 | Degenerate (early) | | | | | | |

[a] Average red fluorescence intensity relative to *mCherry54*.
Average highest fluorescence values (*$CGS4^{hypo}$* healthy) were used as cut-off for thresholding fetal chimaerism (bold).
n, number of fetuses; WT, wild-type.

**Table S12.** Summary of different chimaerism assays for TKO↔*mCherry* aggregation fetuses.

| Cell line | Strain | Fetus ID | Health status | mCherry[a] | | Endpoint PCR | | | | ddPCR (mCherry) | | | | Category |
|---|---|---|---|---|---|---|---|---|---|---|---|---|---|---|
| | | | | Fetus | Kidney | Heart | | Kidney | | Brain | Heart | Kidney | Liver | |
| | | | | | | OFF3 | OFF4 | OFF3 | OFF4 | | | | | |
| OFF3 (male) ↔ OFF4 (female) | *CGS4*$^{hypo}$ ↔ *mCherry38* | F04 | Healthy | **0.07** | **0.41 ± 0.02** | + | - | + | - | ND | ND | 0.01 ± 0.00 | 0.00 ± 0.00 | Chimaera |
| | | F05 | Healthy | 0.05 | 0.06 ± 0.00 | + | - | + | - | ND | ND | 0.00 ± 0.00 | 0.00 ± 0.00 | Host (OFF3) |
| | | F06 | Healthy | **2.05** | **2.40 ± 0.93** | + | + | + | + | 13.09 ± 0.53 | 0.88 ± 0.05 | 1.00 ± 0.08 | 6.19 ± 0.45 | Chimaera |
| | | F09 | Healthy | 0.05 | **0.15 ± 0.03** | + | - | + | - | ND | ND | ND | ND | Chimaera |
| | | F10 | Healthy | 0.05 | **0.20 ± 0.00** | + | - | + | - | ND | 0.00 ± 0.00 | ND | ND | Chimaera |
| OFF3 (male) ↔ OFF4 (female) | *CGS4*$^{hypo}$ ↔ *mCherry54* | F22 | Healthy | 0.01 | 0.02 ± 0.00 | + | - | + | - | ND | ND | ND | ND | Host (OFF3) |
| | | F23 | Healthy | 0.02 | **0.15 ± 0.00** | + | - | + | - | ND | ND | ND | ND | Chimaera |
| | | F24 | Degenerate (late) | 0.02 | NS | + | - | + | - | NS | ND | NS | NS | Host (OFF3) |
| | | F27 | Degenerate (late) | **0.06** | NS | + | + | + | + | NS | 2.47 ± 0.31 | NS | NS | Chimaera |
| | | F31 | Healthy | **0.21** | **0.42 ± 0.01** | + | + | + | + | 0.83 ± 0.14 | 0.34 ± 0.03 | 0.38 ± 0.03 | 0.36 ± 0.02 | Chimaera |
| | | F32 | Healthy | **0.66** | **0.84 ± 0.12** | + | + | + | + | 2.18 ± 0.10 | 3.51 ± 0.15 | 1.75 ± 0.09 | 2.55 ± 0.11 | Chimaera |
| | | F33 | Degenerate (early) | **0.19** | **0.25 ± 0.01** | - | - | + | + | 2.54 ± 0.12 | 2.75 ± 0.13 | 4.83 ± 0.23 | NS | Donor (OFF4) |
| | | F34 | Healthy | **0.82** | **1.20 ± 0.00** | - | - | + | + | 1.84 ± 0.11 | 3.87 ± 0.16 | 2.33 ± 0.13 | 2.72 ± 0.13 | Donor (OFF4) |
| | | F35 | Hydrops | **0.19** | 0.06 ± 0.01 | + | + | + | + | 0.66 ± 0.11 | 0.62 ± 0.04 | 0.29 ± 0.04 | 0.84 ± 0.05 | Chimaera |
| | | F36 | Healthy | **0.18** | **0.26 ± 0.02** | + | + | + | + | 0.28 ± 0.04 | 1.01 ± 0.05 | 0.35 ± 0.04 | 1.43 ± 0.08 | Chimaera |
| | | F37 | Healthy | **0.76** | **0.96 ± 0.12** | - | - | + | + | 2.31 ± 0.26 | 2.48 ± 0.12 | 1.90 ± 0.12 | 3.57 ± 0.17 | Donor (OFF4) |
| | | F38 A | Degenerate (late) | **0.09** | NS | - | - | + | + | NS | 4.39 ± 0.41 | NS | NS | Donor (OFF4) |
| | | F38 B | Degenerate (late) | **0.10** | NS | - | - | + | + | NS | 5.26 ± 0.45 | NS | NS | Donor (OFF4) |
| | | F39 | Healthy | **0.65** | **0.79 ± 0.03** | - | - | + | + | 2.84 ± 0.12 | 2.83 ± 0.13 | 1.99 ± 0.13 | 3.24 ± 0.16 | Donor (OFF4) |
| OFF3 (male) ↔ OFF4 (female) | *CGS20*$^{null}$ ↔ *mCherry54* | F25 | Degenerate (early) | **0.08** | **0.13 ± 0.01** | - | - | + | + | 2.89 ± 0.12 | 4.47 ± 0.22 | 2.18 ± 0.12 | NS | Donor (OFF4) |
| | | F26 | Degenerate (early) | **0.07** | 0.03 ± 0.00 | + | + | + | + | 0.98 ± 0.05 | 2.59 ± 0.20 | 3.89 ± 0.27 | NS | Chimaera |
| | | F28 | Healthy | **0.71** | **0.96 ± 0.00** | + | + | + | + | 2.40 ± 0.13 | 3.22 ± 0.15 | 2.08 ± 0.11 | 2.26 ± 0.12 | Chimaera |
| | | F29 | Healthy | **0.39** | **0.57 ± 0.13** | + | + | + | + | 1.17 ± 0.11 | 2.02 ± 0.10 | 1.57 ± 0.08 | 1.92 ± 0.13 | Chimaera |
| | | F30 | Degenerate (late) | **0.08** | NS | - | - | + | + | NS | 6.13 ± 0.36 | NS | NS | Donor (OFF4) |

[a] Average red fluorescence intensity relative to *mCherry54*. Values above wild-type background (0.06 and 0.13 for fetus and kidney, respectively) are in bold.
ND, no mCherry detected; NS, not sampled due to degeneration.

### Supplementary Figures

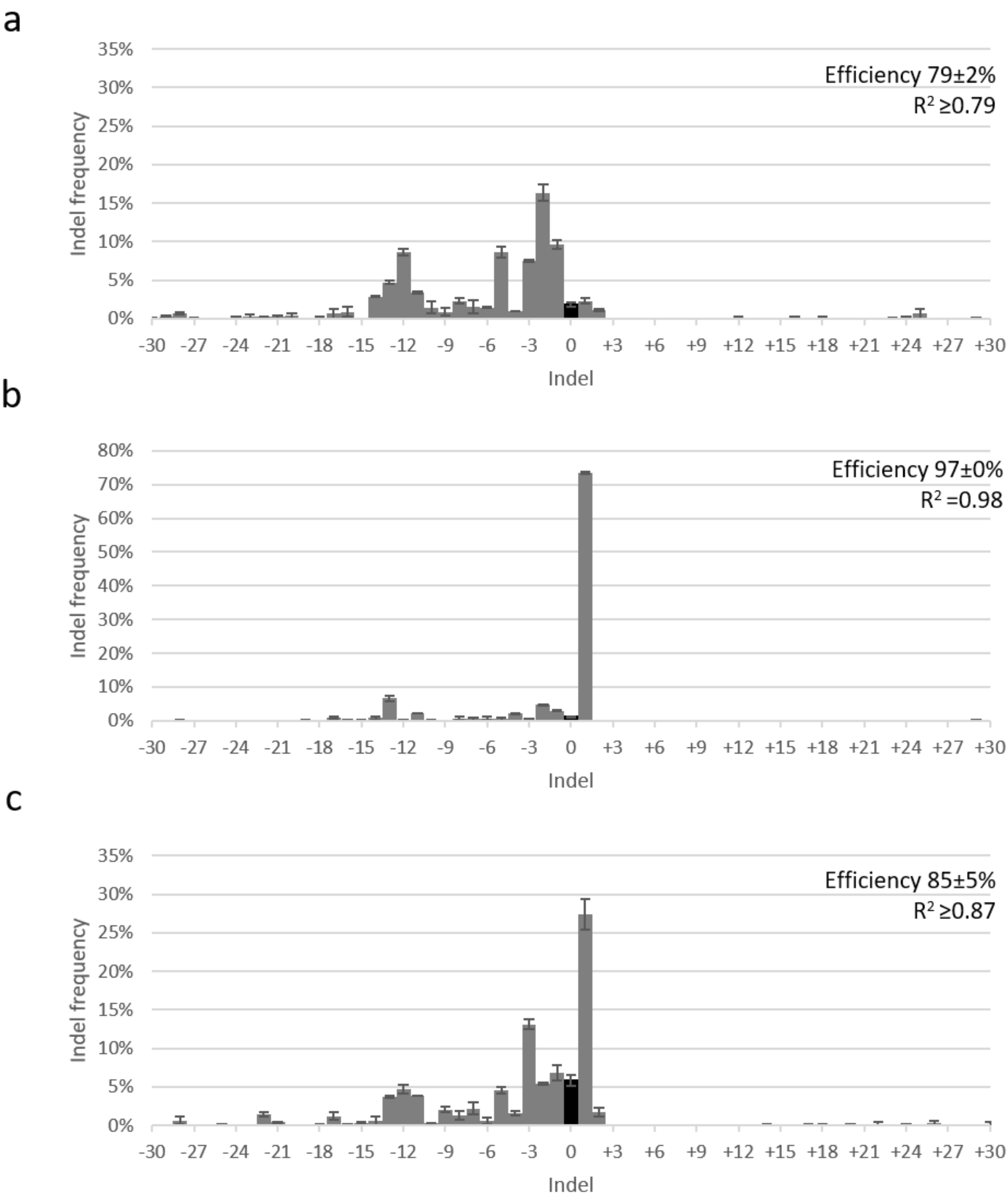


**Fig. S1. Sequence analysis of editing for *SALL1* (a) gRNA1, (b) gRNA2, and (c) gRNA3.** Sanger sequencing reads of edited population from each transfection was run through TIDE to estimate proportion of indels present. Non-edited sequences shown in black. Error bars = SEM, n= 2.

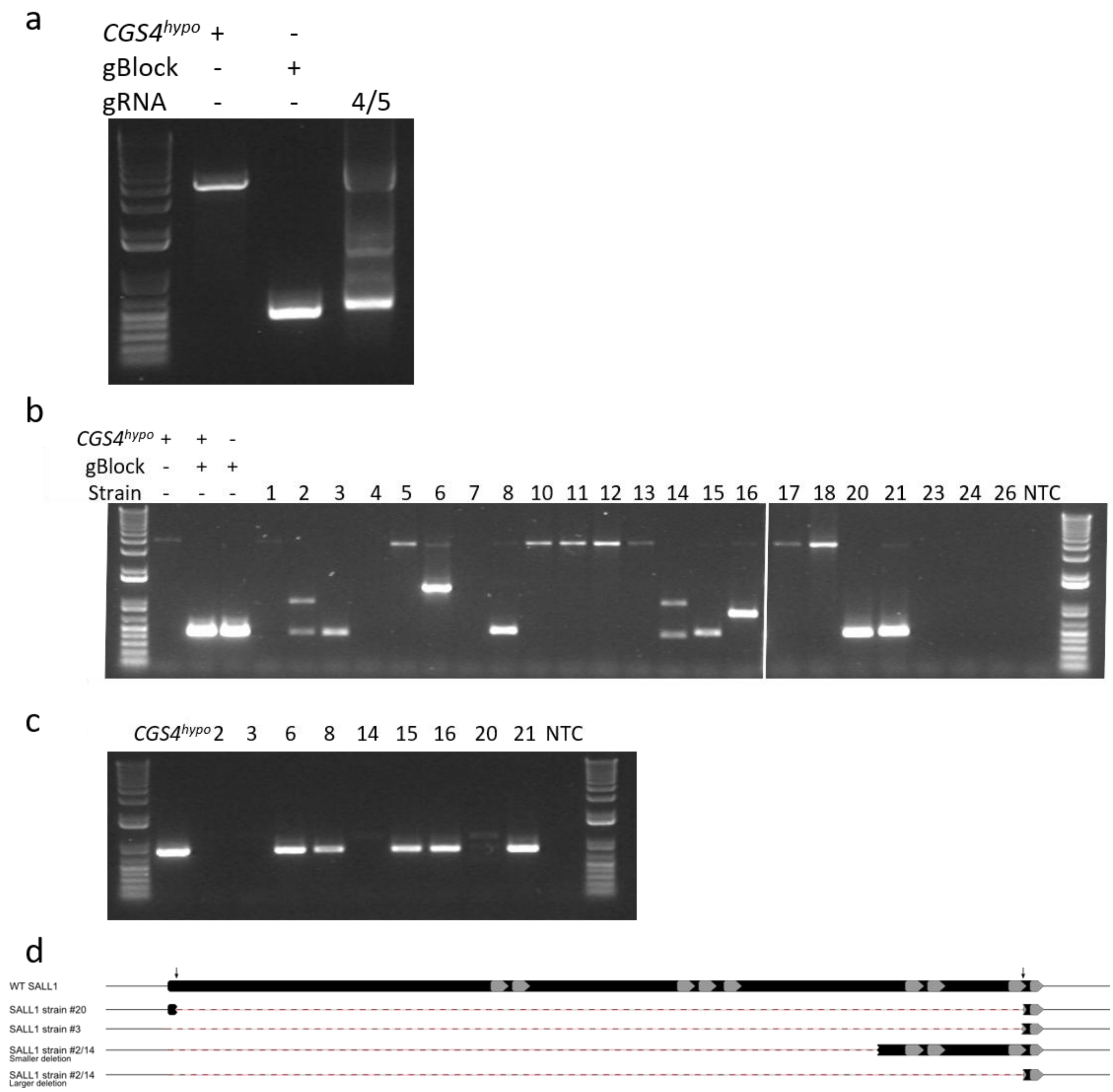


**Fig. S2. Screening of dual guide-edited cells for large deletions.** (a) PCR following transfection of dual guides (gRNA4/5). Parental OFF3 *CGS4*$^{hypo}$ and synthetic, double-stranded DNA fragments to mimic the large deletion strategy (gBlock), generating 3.7 kb amplicon and 0.4 kb amplicons, respectively. (b) PCR assays to identify large deletion in strains isolated from gRNA4/gRNA5 transfection. End-point reaction designed to amplify a 3.7 kb fragment for *CGS4*$^{hypo}$ and 0.4 kb for large deletion gBlock. Control samples containing *CGS4*$^{hypo}$ and gBlocks were also included. All reactions contained a no template control (NTC). Large deletions were predicted in samples that had 1 or 2 bands <1.5 kb. (c) For strains with a predicted large deletion, another PCR was carried out with primers that bound within the intended deleted sequence to confirm predicted editing profile. Positive genomic control *CGS4*$^{hypo}$ and NTC included. (d) Gene structure of the four strains identified with large deletions. Features shown = gRNA binding site (black arrow), CDS (black), functional domains (grey), and deleted areas (dashed red).

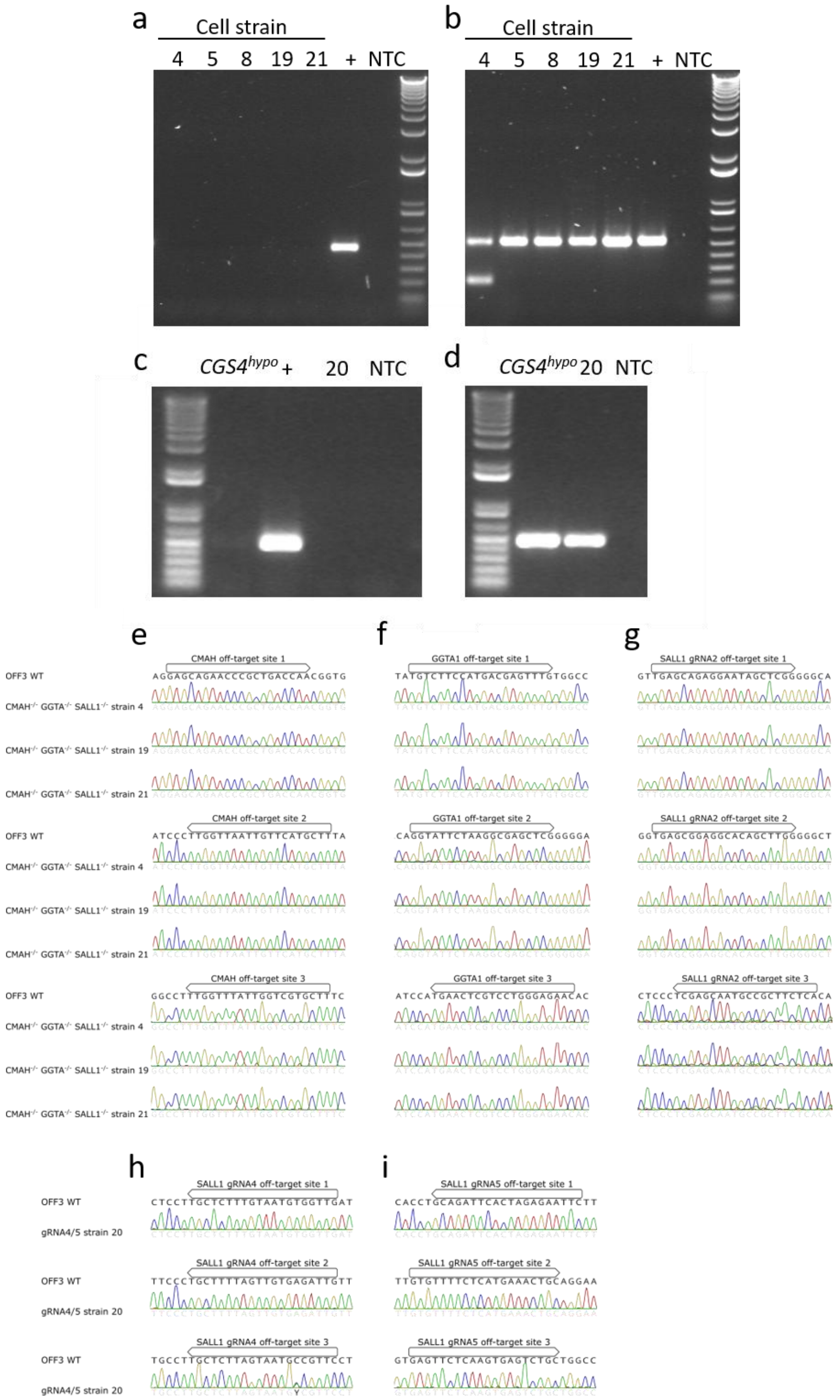


**Fig. S3. PCR screening for plasmid insertion and off-target editing.** (a) *Cas9* plasmid insertion and (b) *GGTA1* genomic DNA control in triple edited, small indel strains. (c) Cas9 plasmid insertion and (d) gRNA4 off-target site 1 genomic DNA control for chosen large deletion strain (20). End-point PCR performed with primers designed to *Cas9* sequence in plasmid backbone. Cas9 and genomic DNA positive control (+) from a OFF3 *CMAH*$^{-/-}$ *GGTA1*$^{-/-}$ confirmed *Cas9* integration strain, genomic DNA control *CGS4*$^{hypo}$ (*Cas9* negative), and no template control (NTC). Small indel strains were screened for off-target sites of (e) *CMAH* gRNA1, (f) *GGTA1*, and (g) *SALL1* gRNA2. Large deletion strains were screened for off-target sites of (h) gRNA4 and (i) gRNA5. Sanger sequencing of regions identified by CRISPOR as most likely off-target sites. Boxes indicate off-target sequence (not including PAM site).

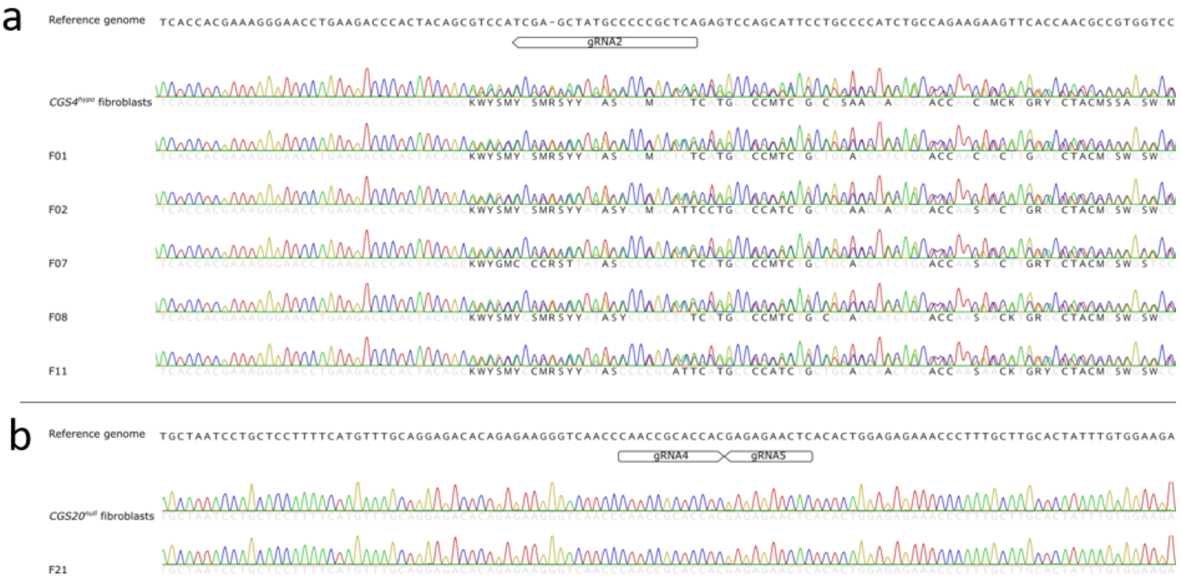


**Fig. S4. Genotyping edited fetuses by Sanger sequencing.** Fetuses sequenced for *SALL1* in (a) single and (b) dual guide editing strategies.

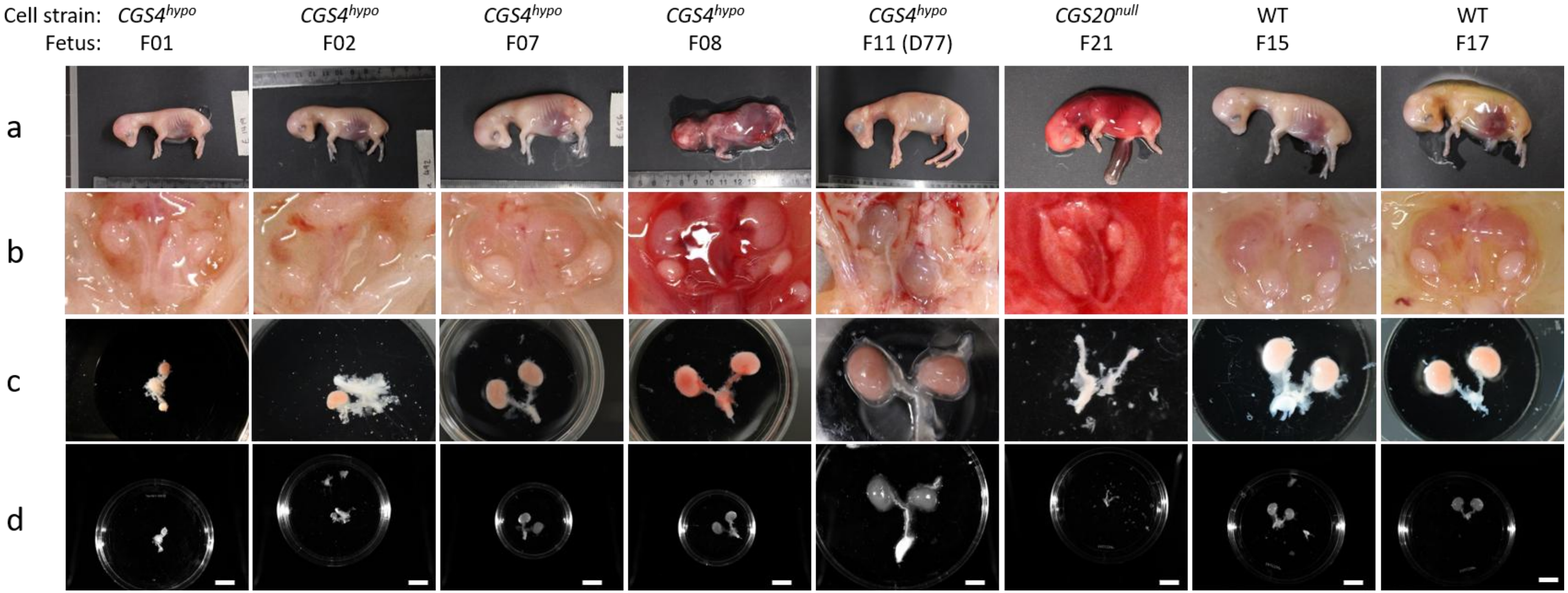


**Fig. S5. Morphology of *SALL1*$^{-/-}$ fetuses on D48 or D77.** Images show (a) whole fetus, (b) *in situ* presentation of kidneys, (c) separated metanephros, and (d) metanephric kidneys at the same scale (Colorimetric image on ChemiDoc). Scale bar = 10 mm.

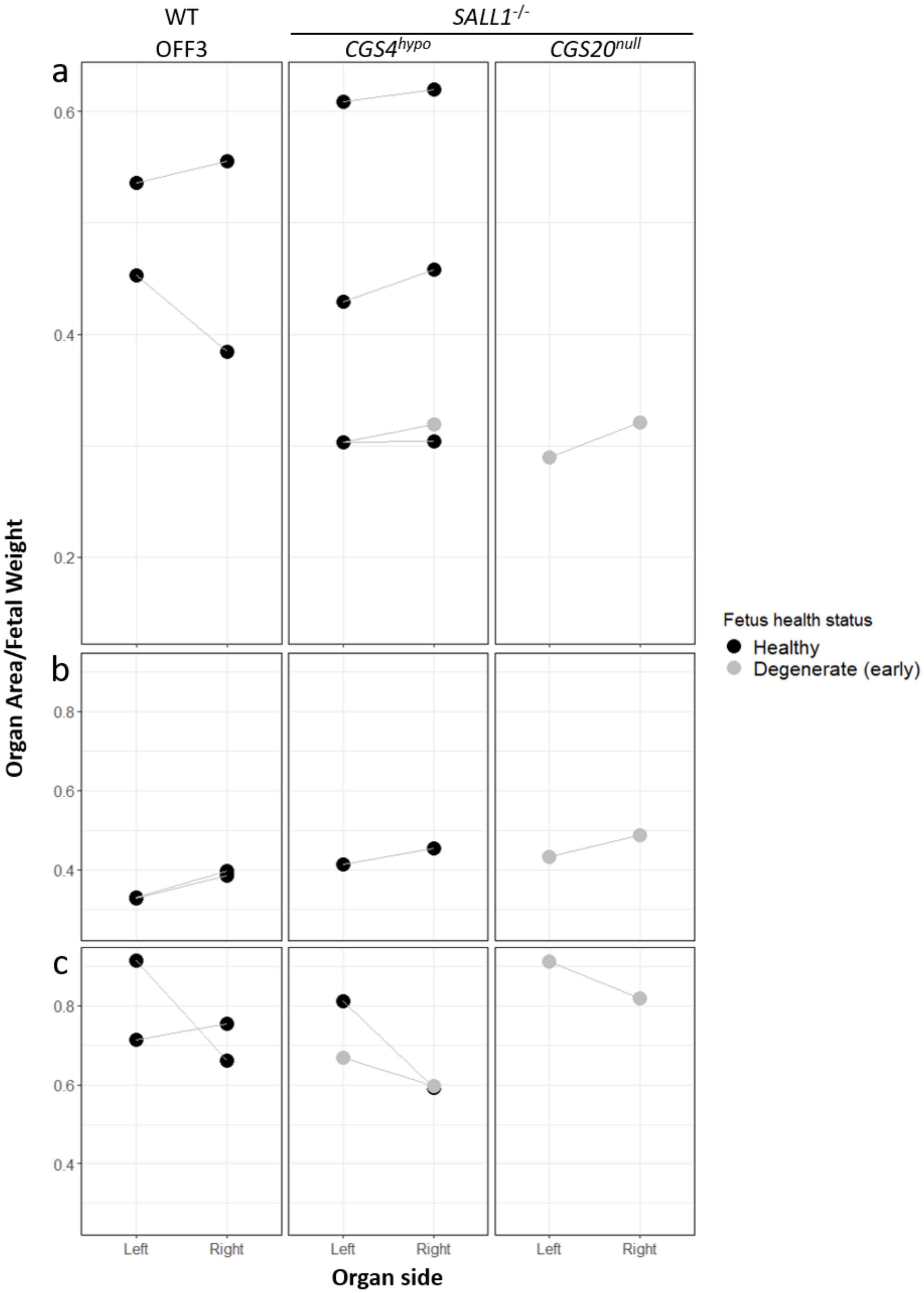


**Fig. S6. Comparison of (a) gonad, (b) adrenal, and (c) mesonephros size in *SALL1*$^{-/-}$ fetuses.** Organ area ($mm^2$) normalized on fetal weight (g). *SALL1* KO fetuses from strain *CGS4*$^{hypo}$ and *CGS20*$^{null}$. WT control males (OFF3). Paired organs from a single fetus are connected with a grey line.

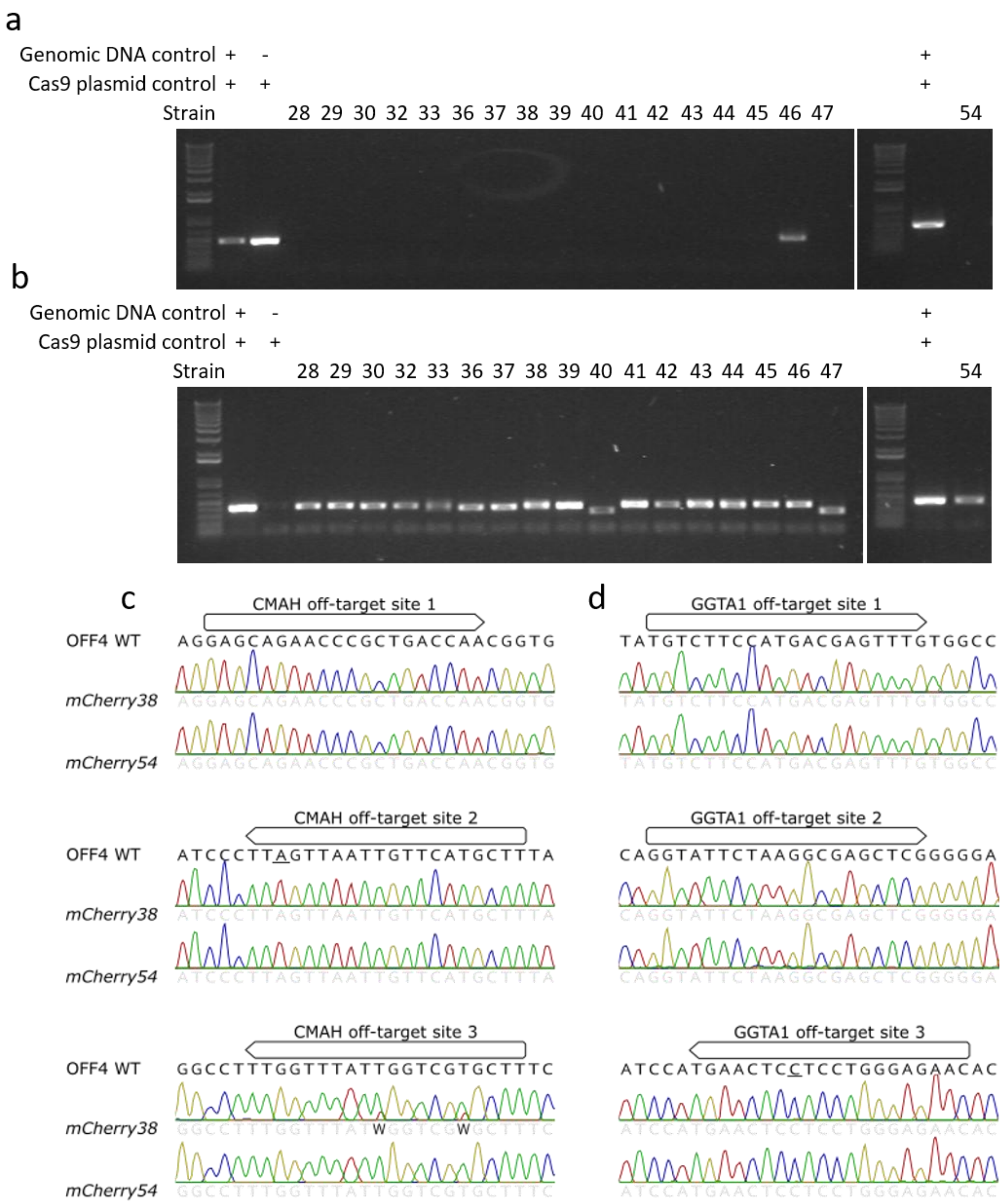


**Fig. S7. PCR screening for plasmid insertion and off-target editing in *mCherry* donor strains.** (a) *Cas9* plasmid insertion and (b) *CMAH* DNA control in *CMAH/GGTA1*-edited *mCherry* strains. End-point PCR performed with primers designed against *Cas9* sequence in plasmid backbone. Positive controls (+) include OFF3 *CMAH*$^{-/-}$ *GGTA1*$^{-/-}$ 41 (confirmed *Cas9* integration strain) and *Cas9* plasmid control (PX459). Off-target screen of top three sites for (c) *CMAH* and (d) *GGTA1* in selected strains. Sanger sequencing of regions identified by CRISPOR as most likely off-target sites. Boxes indicate off-target sequence (not including PAM site). *CMAH* off-target site 2 and *GGTA1* off-target site 3 have a SNP in the WT sequence of OFF4 (underlined) compared to reference genome.

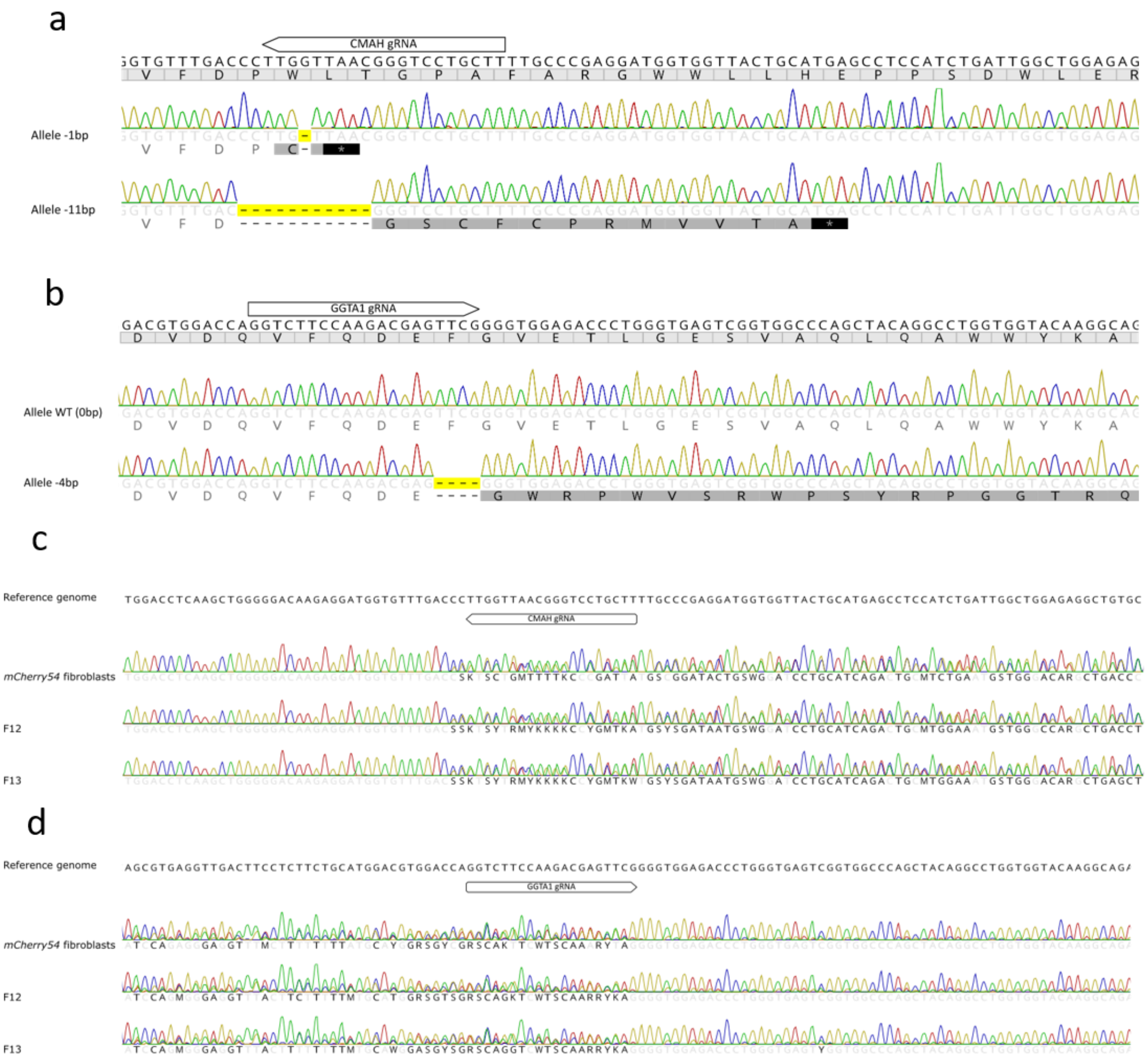


**Fig. S8. Sanger sequences of cloned mCherry fetuses.** *mCherry54* sequenced for (a) *CMAH* and (b) *GGTA1*. Translation shown under nucleotide sequence. PCR products that could not be separated on agarose gel were sub-cloned into bacteria to sequence individual alleles. Edited nucleotides highlighted yellow (with changes to amino acid sequence shown underneath in black within shaded boxes). Matching nucleotides/amino acids in light grey font. Black boxes containing star indicate stop codon. *mCherry54* fetuses sequenced for (c) *CMAH* and (d) *GGTA1*.

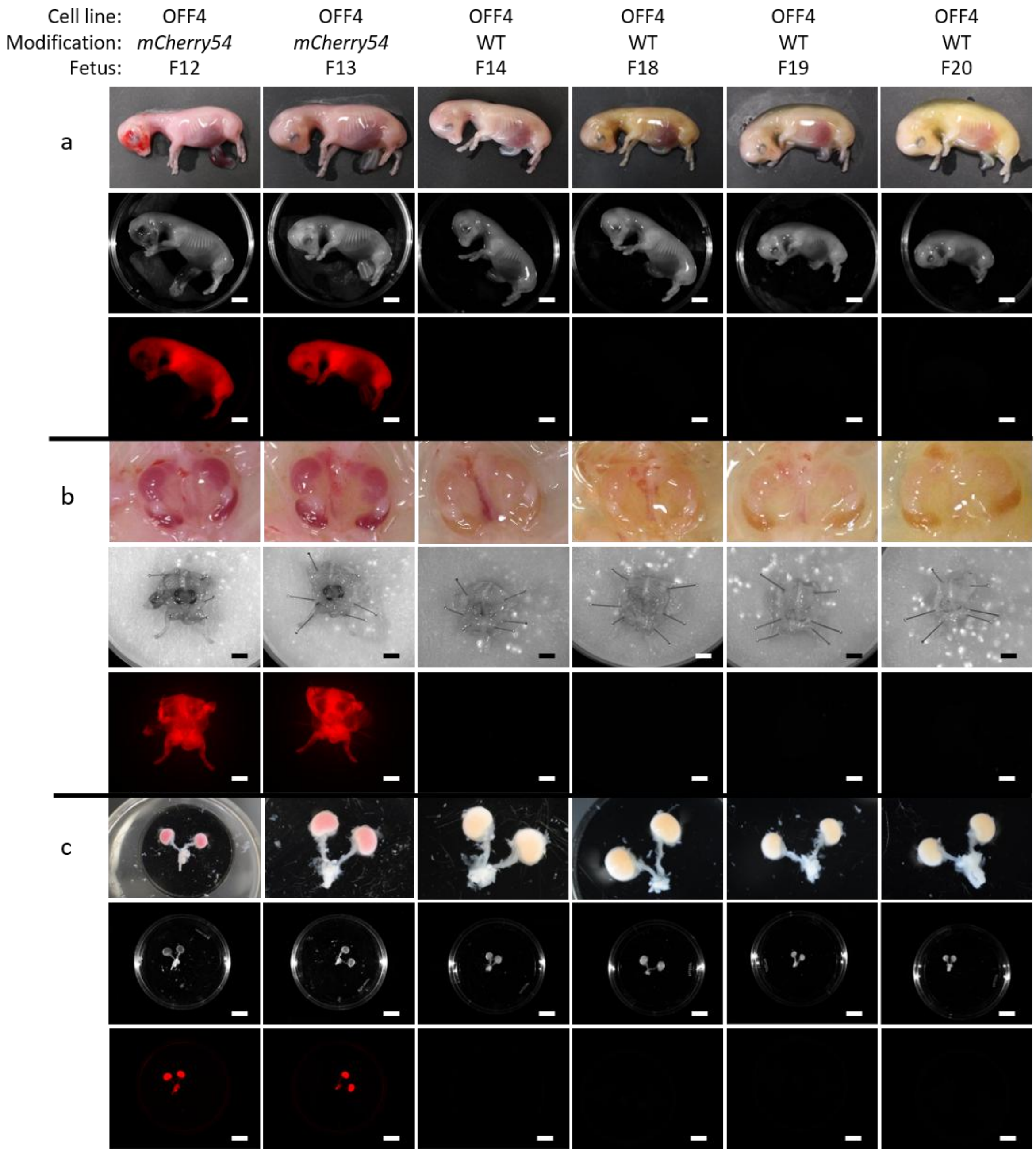


**Fig. S9. Morphology of healthy mCherry fetuses on D48.** Images show (a) whole fetus, (b) in situ presentation of kidneys, and (c) separated metanephros. Within each panel, the top image is taken by digital single-lens reflex camera, middle by ChemiDoc (colorimetric), and bottom by ChemiDoc (0.2 s exposure on DyLight550 channel). Control fetuses derived from OFF4 WT parental line without edits or insertions. Scale bar = 10 mm.

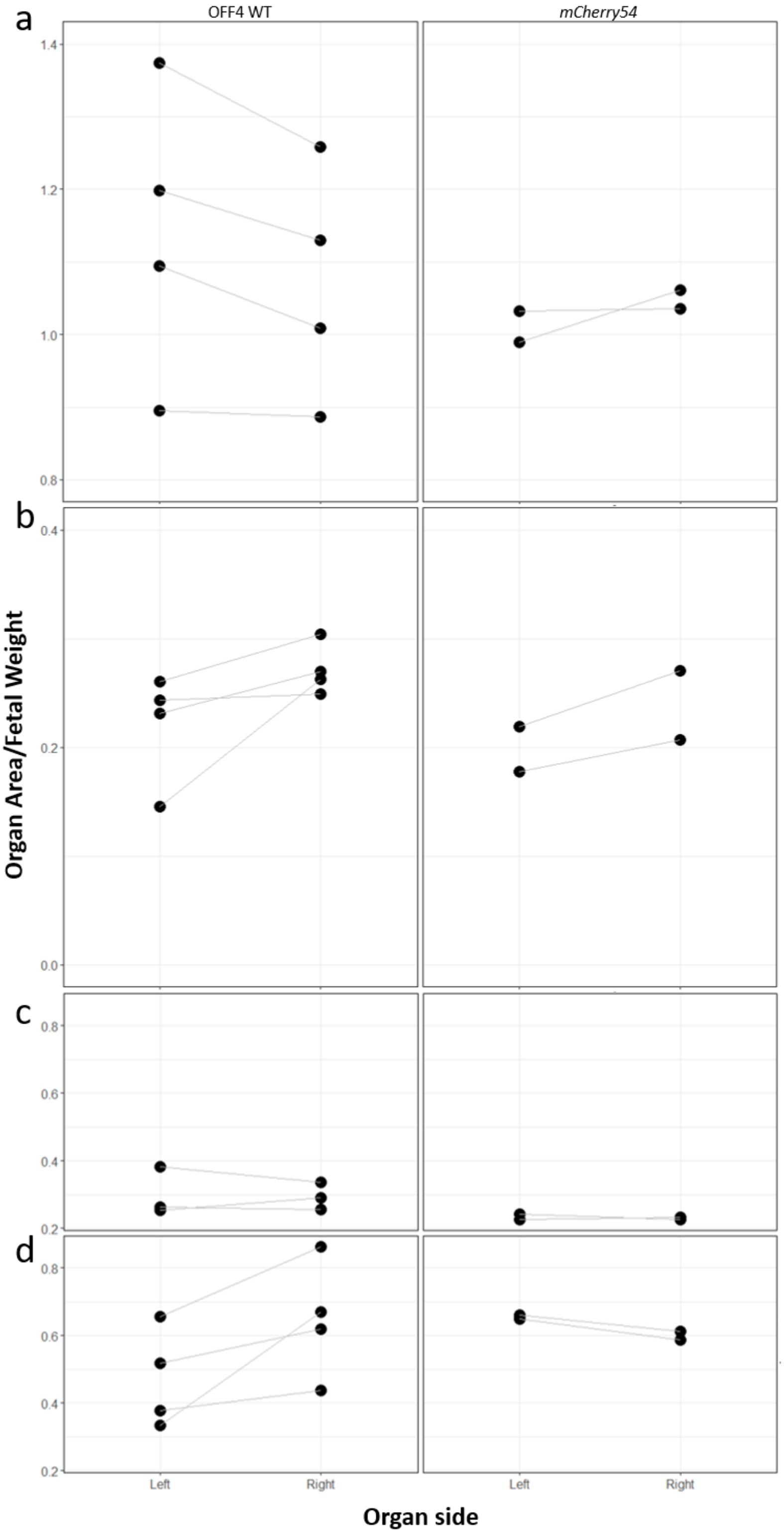


**Fig. S10. Organ sizes in recovered healthy mCherry fetuses on D48.** Comparison of (a) metanephros, (b) gonad, (c) adrenal, and (d) mesonephros in collected healthy fetuses. Organ area ($mm^2$) normalized on fetal weight (g). mCherry fetuses only from strain 54 as no fetuses survived to D48 from strain 38. WT controls derived from OFF4 female line. Paired organs from a single fetus are connected with a grey line.

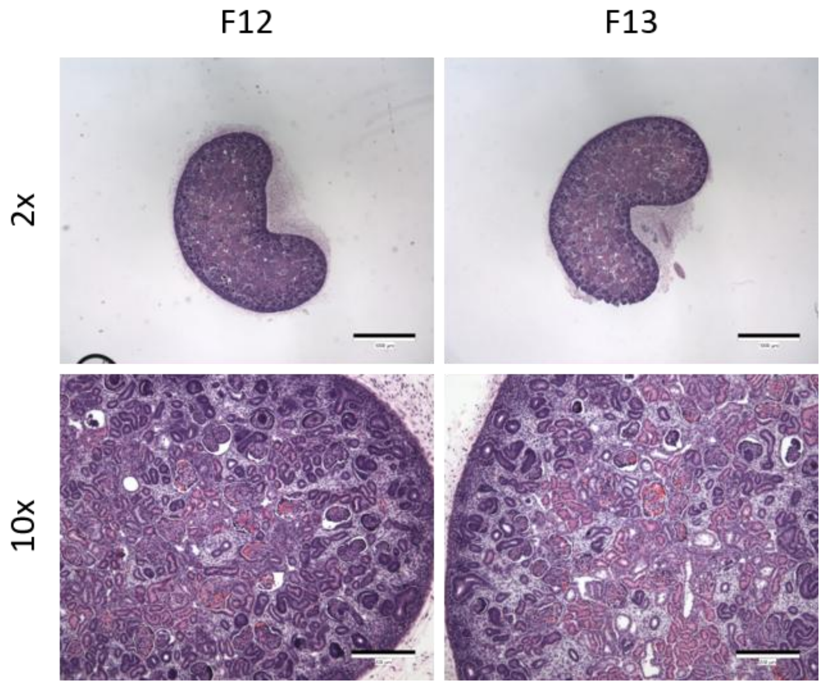


**Fig. S11. Histology of *mCherry54* metanephric kidneys.** Hematoxylin/eosin-stained sections imaged with 2x objective (scale bar = 1000 µm), and magnified view imaged with 10x objective (scale bar = 200 µm).

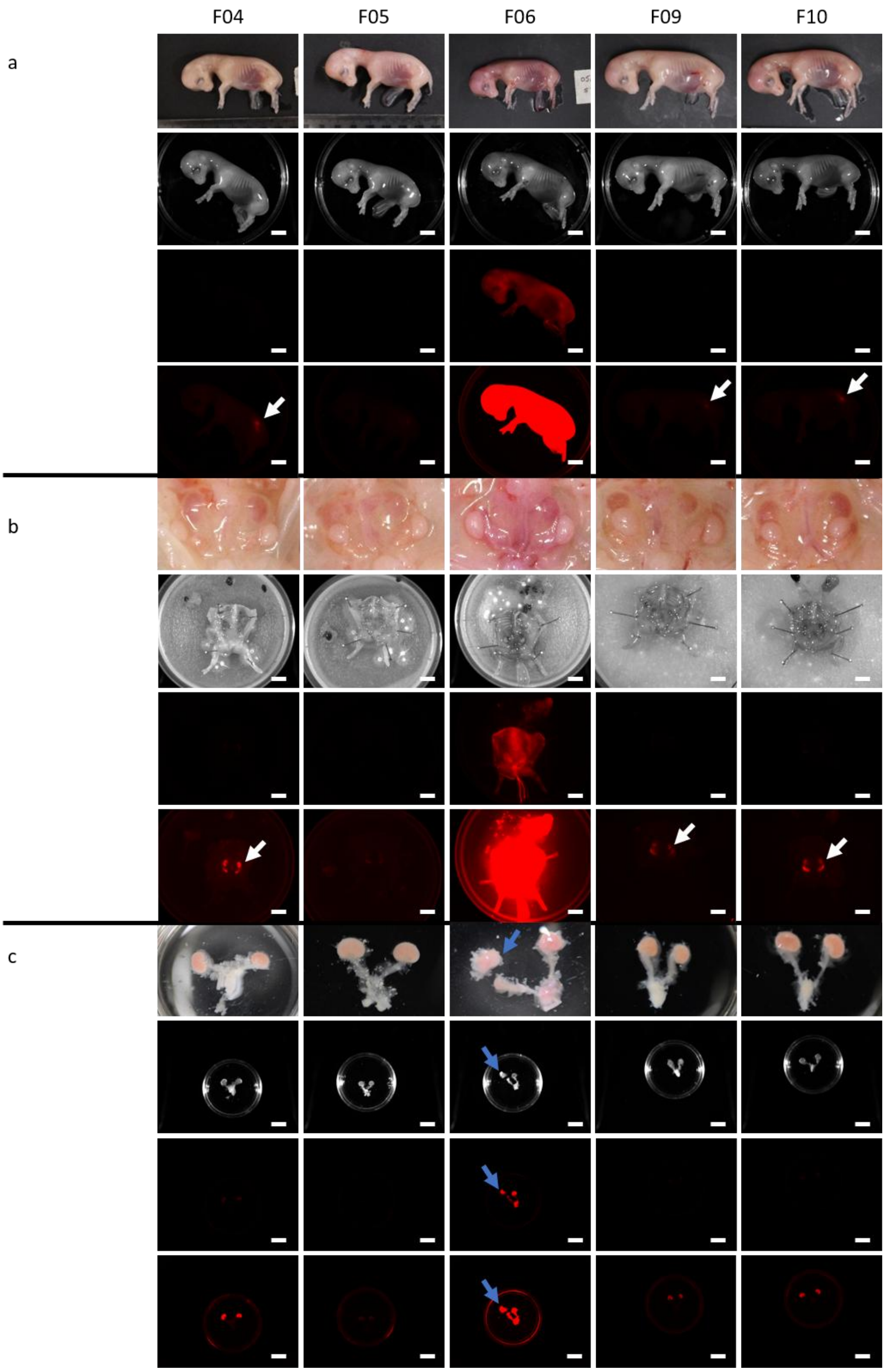


**Fig. S12.** ***CGS4$^{hypo}$↔mCherry38*** **aggregation fetuses on D48.** Images show (a) whole fetus, (b) *in situ* presentation of kidneys, and (c) separated metanephros. Within each panel, the top image is taken by digital single-lens reflex camera, second by ChemiDoc (colorimetric), and third and fourth by ChemiDoc (0.2 s and 2s exposure, respectively, on DyLight550 channel). White arrows indicate fluorescent metanephric and mesonephric kidneys visible with 2s exposure through body wall in fetal image or *in situ* following dissection. One adrenal was left in the Petri dish after separation from smaller metanephric kidney for F06 (blue arrow) Scale bar = 10 mm.

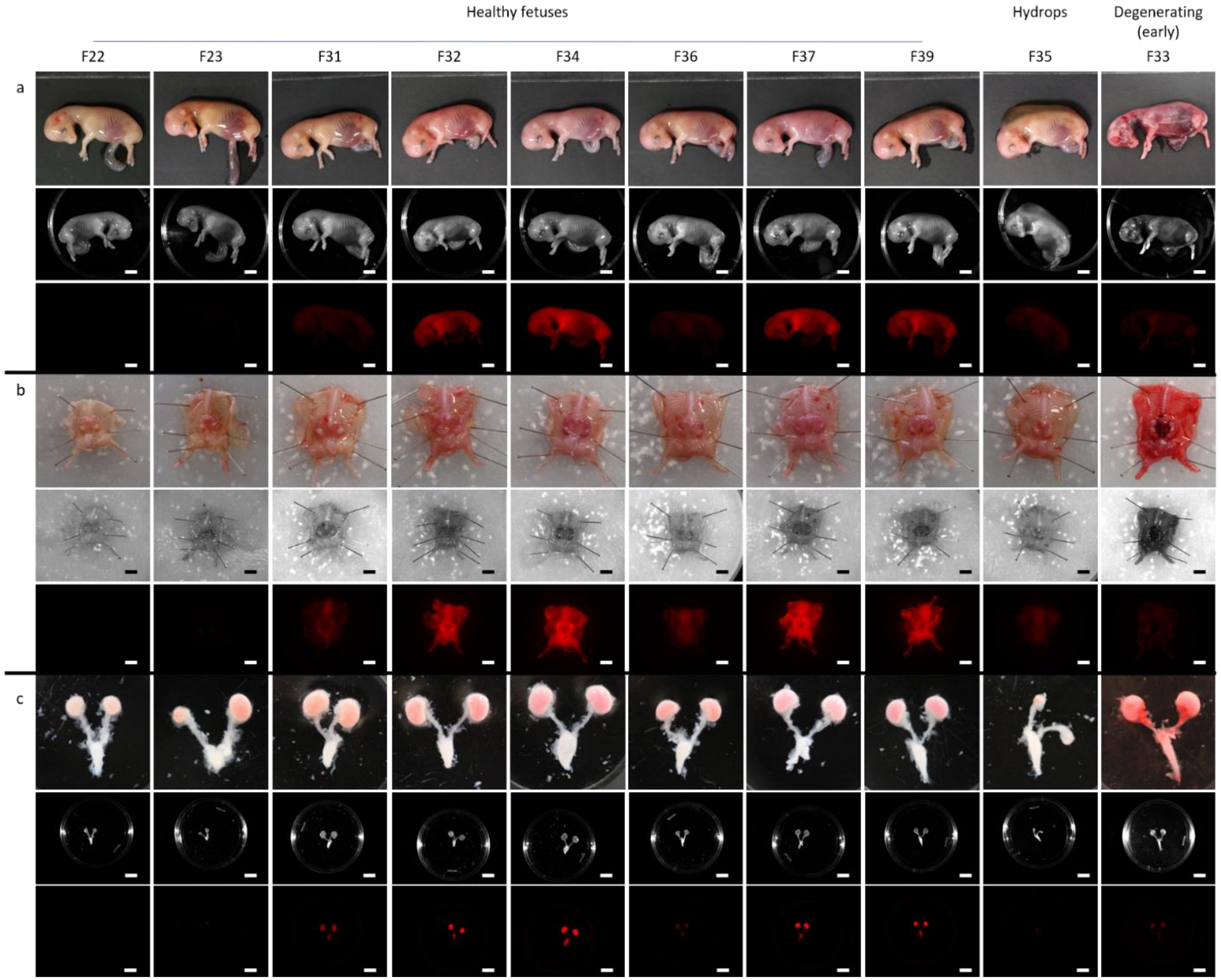


**Fig. S13.** ***CGS4$^{hypo}$↔mCherry54*** **aggregation fetuses on D48.** Images show (a) whole fetus, (b) *in situ* presentation of kidneys, and (c) separated metanephros. Within each panel, the top image is taken by digital single-lens reflex camera, middle by ChemiDoc (colorimetric), and bottom by ChemiDoc (0.2 s exposure on DyLight550 channel). Scale bar = 10 mm.

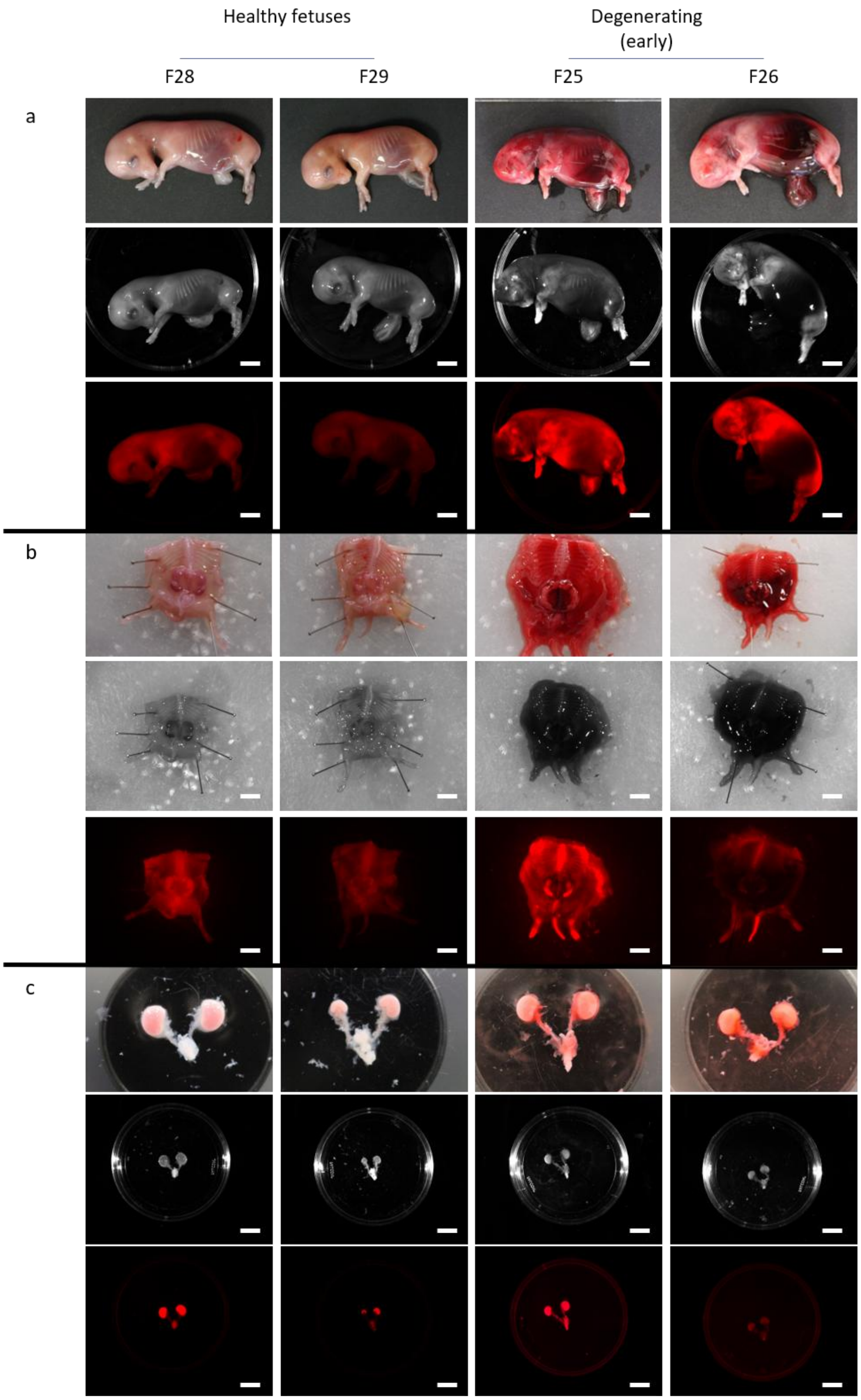


**Fig. S14. *CGS20$^{null}$↔mCherry54* aggregation fetuses on D48.** Images show (a) whole fetus, (b) *in situ* presentation of kidneys, and (c) separated metanephros. Within each panel, the top image is taken by digital single-lens reflex camera, middle by ChemiDoc (colorimetric), and bottom by ChemiDoc at 0.2s exposure (healthy fetuses) or 2s (early degenerating fetuses). Scale bar = 10 mm.

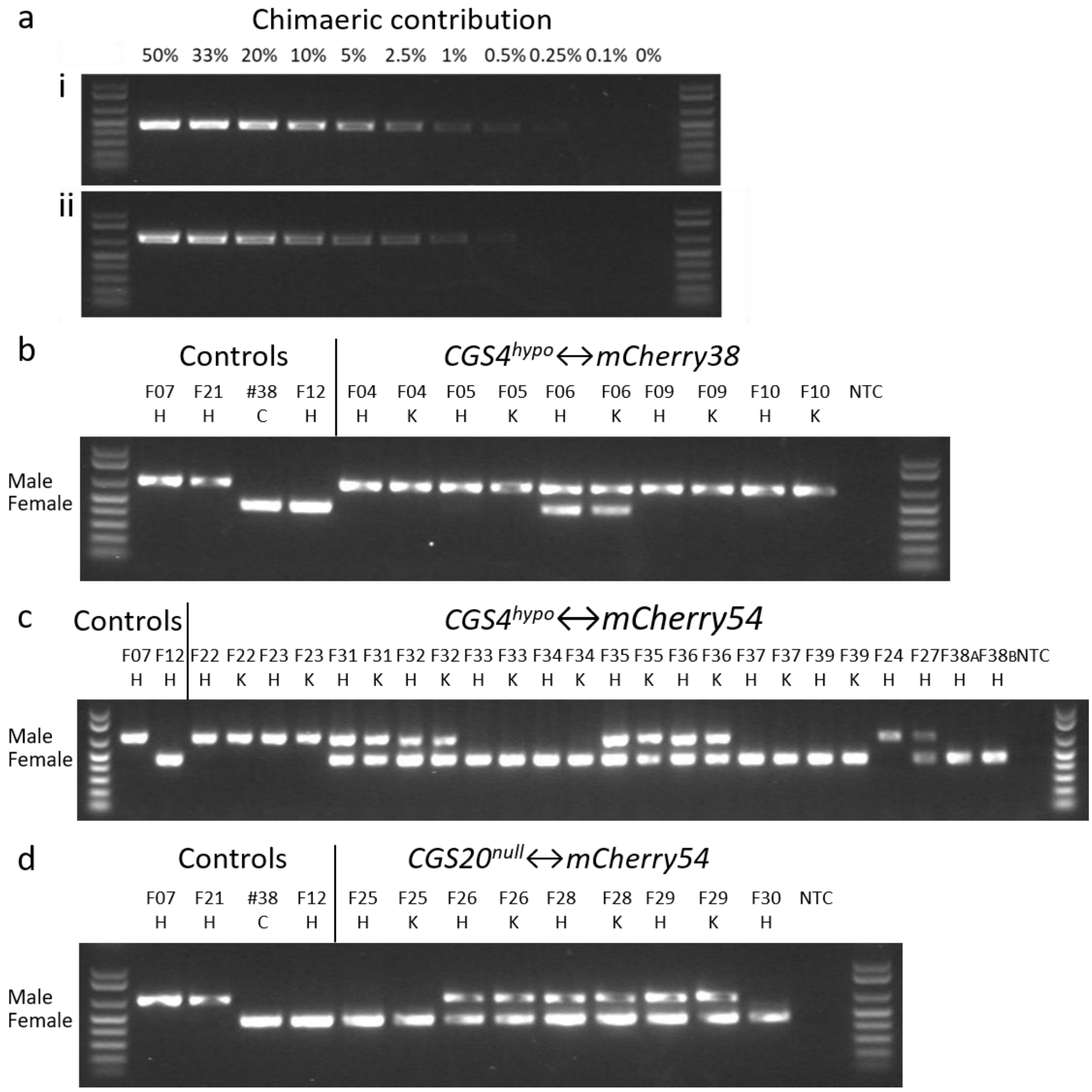


**Fig. S15. Endpoint PCR to determine fetal sex chimaerism.** Fetuses derived from (a) *CGS4*$^{hypo}$↔*mCherry38*, (b) *CGS4*$^{hypo}$↔*mCherry54*, and (c) *CGS20*$^{null}$↔*mCherry54* aggregation. Male primers designed for *DDX3Y* (Y chromosome), and female primers to bind within edited region of *CMAH* in *CGS4*$^{hypo}$ and *CGS20*$^{null}$. DNA was isolated from heart (H), kidney (K), and cultured cells (C). Positive controls included male *CGS4*$^{hypo}$ F07 and *CGS20*$^{null}$ F21, female *mCherry38* cells and *mCherry54* F12, and no template control (NTC). (d) Female DNA assay was repeated with increased DNA concentration (40–80 ng) for chimaeras with fluorescent kidneys but no signal in endpoint PCR. Female-specific *CMAH* assay performed alongside genomic *CMAH* DNA control. Chimaeric *CGS4*$^{hypo}$↔*mCherry54* fetus (F31) served as a positive control. Negative controls included non-fluorescent fetus (F05) and NTC.

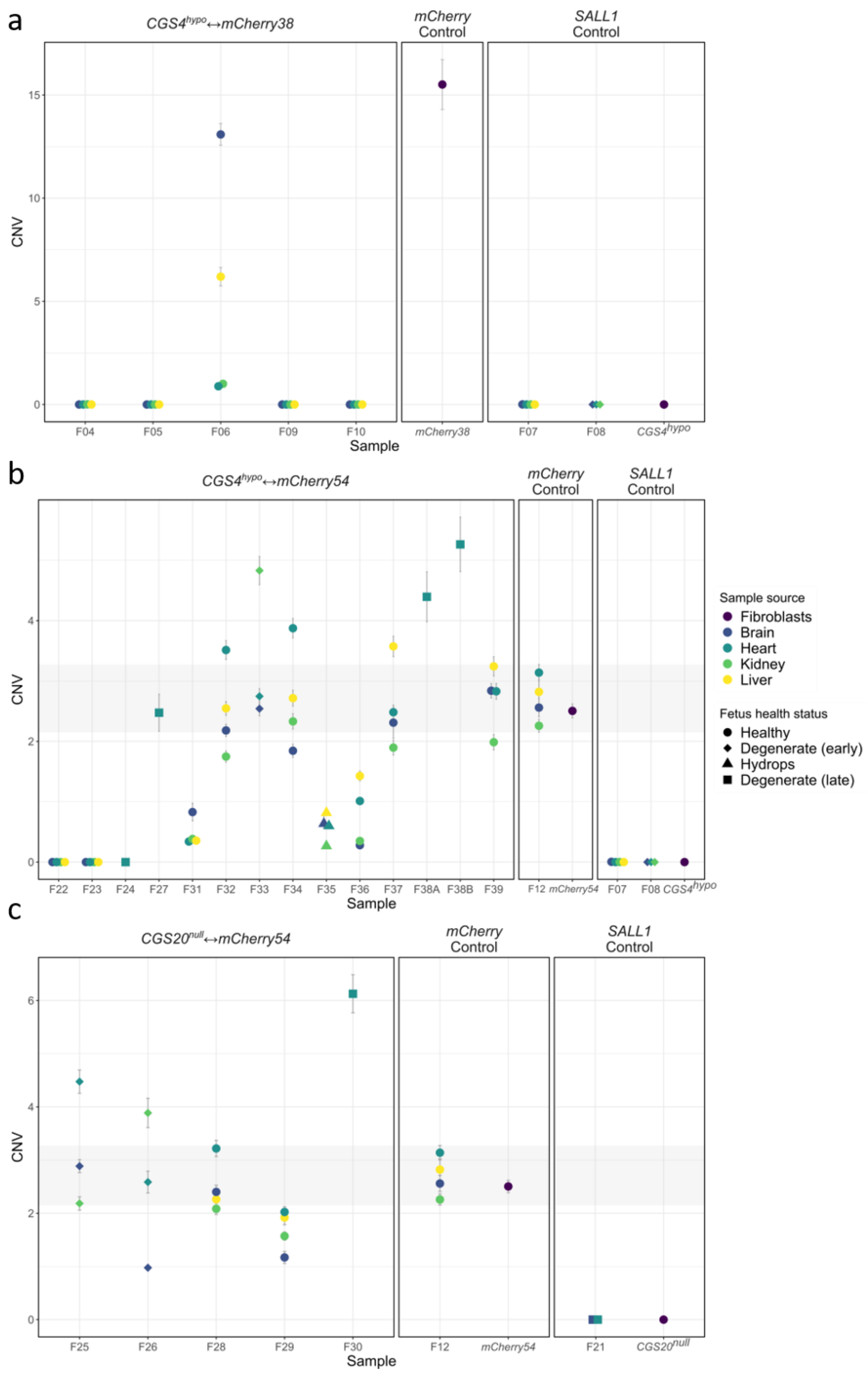


**Fig. S16. ddPCR to determine fetal mCherry chimaerism.** Fetuses derived from (a) *CGS4*$^{hypo}$↔*mCherry38*, (b) *CGS4*$^{hypo}$↔*mCherry54*, and (c) *CGS20*$^{null}$↔*mCherry54*. DNA isolated from fetuses and copies of *mCherry* normalized on *CSN2* reference. Fetal health status shown as data point shape (circle = healthy, diamond = early-stage degeneration, triangle = hydrops, and square = late stage degeneration). Color indicates source of the DNA sample (cultured fibroblasts = dark purple, brain = dark blue, heart = teal, kidney = green, and liver = yellow). Control tissue samples for *CGS4*$^{hypo}$ were from fetuses F07 and F08, and for *CGS20*$^{null}$ from F21. Control *mCherry54* fetal samples were from F12. No fetuses from *mCherry38* survived to D48, so only an original edited cell sample control was included for this strain. Grey shading indicates range observed by *mCherry54* control fetus (F12).

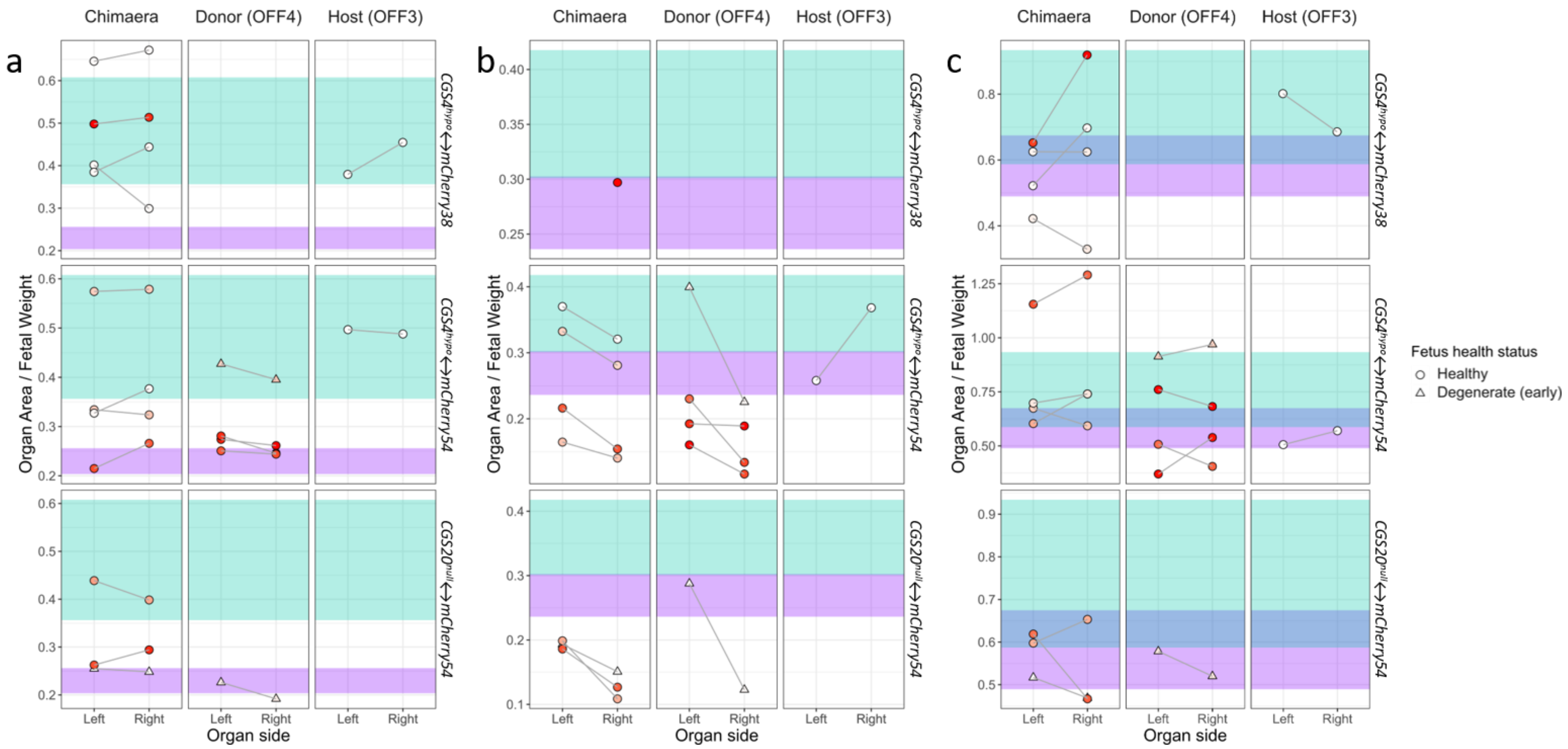


**Fig. S17. Comparison of (a) gonad, (b) adrenal, and (c) mesonephros size in chimaeric fetuses.** Normalized organ area from collected healthy *CGS4*$^{hypo}$*↔mCherry38*, *CGS4*$^{hypo}$*↔mCherry54*, and *CGS20*$^{null}$*↔mCherry54* fetuses. mCherry fluorescence level indicated by red coloring of circles. 95% CI for WT female (lavender) and male (aqua). Paired organs from single fetuses are connected with a grey line.

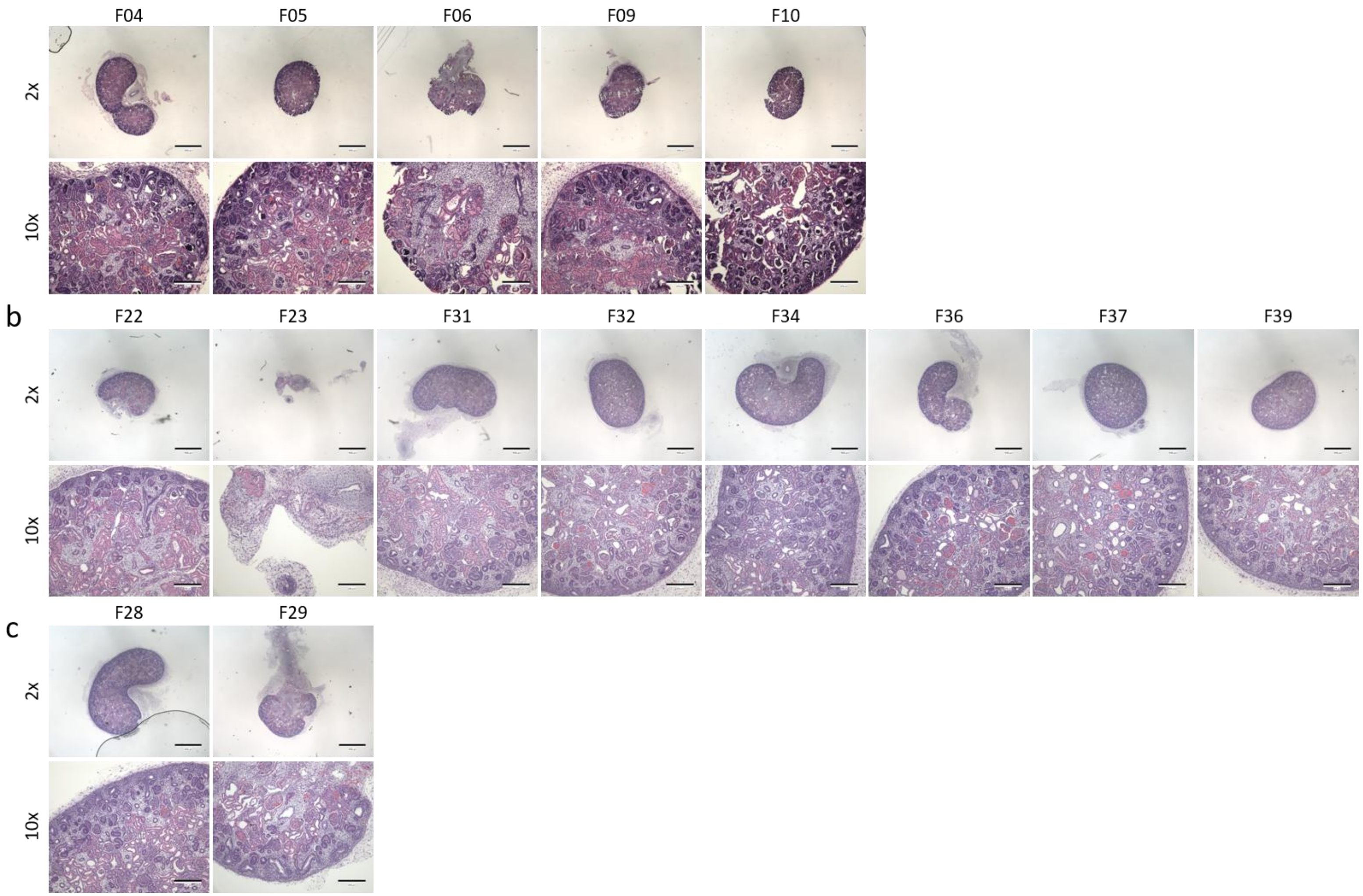


**Fig. S18. Histology of chimaeric metanephric kidneys.** Hematoxylin/eosin-stained sections from (a) *CGS4*$^{hypo}$↔*mCherry38*, (b) *CGS4*$^{hypo}$↔*mCherry54*, and (c) *CGS20*$^{null}$↔*mCherry54* chimaeras imaged with 2x objective (scale bar = 1000 µm), and magnified view imaged with 10x objective (scale bar = 200 µm).

### Supplementary Methods

#### PX459-Hygro production

Puromycin resistance was removed from the PX459 plasmid by digestion using *EcoR*I sites identified 5' of the cleavage peptide and 3' of the puromycin resistance sequences. The hygromycin sequence was based off plasmids within the Addgene database that contained both hygromycin resistance and Cas9 enzyme sequences (pCRISPR-S12, Addgene plasmid 84031 (Li et al., 2016); Hygro-Cas9, Addgene plasmid 86883 (Shi et al., 2017); e-ciCas9_pcDNA5, Addgene plasmid 100553 (Rose et al., 2017)), and the nucleotide sequence modified (maintaining amino acid sequence) to remove any internal *EcoR*I sites. The finalized hygromycin sequence (BJO288) was ordered as a GeneArt™ Strings™ DNA Fragment (Invitrogen, Waltham, MA, USA) and resuspended in MilliQ $H_2O$ to 100 ng/µL, as per Invitrogen instructions. A-tailed hygromycin fragments were inserted into pGEM®-T Easy vector (Promega, Madison, WI, USA), according to kit instructions with a 1:1 vector insert ratio (50 ng of vector:18.45 ng of BJO288), and transformed into bacteria. Amplified plasmids were sequence verified before being used to insert the hygromycin sequence into PX459. Hygromycin vector and PX459 (1 µg of DNA total) were digested with 15–20 Units of *EcoR*I-HF (New England Biolabs, Ipswich, MA, USA) in CutSmart buffer (New England Biolabs) to a total reaction volume of 20–50 µL and incubated at 37°C for 20 min or overnight. PX459 digestion also included 1 µL of thermosensitive alkaline phosphatase (TSAP) to remove 5' phosphate groups, preventing plasmid recircularization. Following digestion, PX459 was incubated at 74°C for 15 min to inactivate TSAP. Digested fragments were ligated together at a 1:2 vector insert ratio (35 ng of PX459:9 ng of Hygromycin) using 1 µL of T4 ligase (Promega) in 10 µL of 2X T4 ligase buffer (from pGEM®-T Easy Vector kit) made up to 20 µL with MilliQ $H_2O$. The reaction was incubated for 3 h at room temperature, before transformation into DH5α (Zymo Research, Irvine, CA, USA) or TOP10 *Escherichia coli* (Thermo Fisher, Waltham, MA, USA). The plasmid was amplified and isolated using PureLink™ HiPure Miniprep Kit (Thermo Fisher), following the manufacturer's instructions. A diagnostic restriction enzyme digestion was performed on isolated plasmids to identify if the hygromycin sequence was inserted in the correct orientation. Two *SacII* recognition sites occurred within the final plasmid, one within PX459 and one within the hygromycin sequence. Digestion with *SacII*, therefore, allowed determination of no hygromycin insertion (single 9.5 kb band), wrong orientation (6.7 and 2.8 kb bands), or correct orientation (7.3 and 2.2 kb bands). At least 250 ng of plasmid was digested with 1 µL of *SacII* (New England Biolabs) in CutSmart buffer, with the final reaction made up to 20 µL. Plasmids were incubated at 37°C for 1.5 h or overnight before being run on a 1% agarose gel to confirm banding pattern.

Plasmids identified with insert in correct orientation were then sent for sequencing with primers designed to sequence from within the hygromycin resistance gene into PX459 (GL279) and from PX459 into the hygromycin sequence (GL781). Correctly inserted plasmids were amplified further using PureLink™ HiPure Maxiprep Kit (Thermo Fisher).


Li, L., Gao, F., & Wu, S. (2016). An episomal CRISPR/Cas9 system to derive vector-free gene modified mammalian cells. *Protein & Cell*, *7*(9), 689–691. https://doi.org/10.1007/s13238-016-0299-9

Rose, J. C., Stephany, J. J., Valente, W. J., Trevillian, B. M., Dang, H. V., Bielas, J. H., Maly, D. J., & Fowler, D. M. (2017). Rapidly inducible Cas9 and DSB-ddPCR to probe editing kinetics. *Nature Methods*, *14*(9), 891–896. https://doi.org/10.1038/nmeth.4368

Shi, Z.-D., Lee, K., Yang, D., Amin, S., Verma, N., Li, Q. V., Zhu, Z., Soh, C.-L., Kumar, R., Evans, T., Chen, S., & Huangfu, D. (2017). Genome editing in hPSCs reveals GATA6 haploinsufficiency and a genetic interaction with GATA4 in human pancreatic development. *Cell Stem Cell*, *20*(5), 675-688.e6. https://doi.org/10.1016/j.stem.2017.01.001
